# Cryo-ET of actin cytoskeleton and membrane structure in lamellipodia formation using optogenetics

**DOI:** 10.1101/2024.08.13.607852

**Authors:** Hironori Inaba, Tsuyoshi Imasaki, Kazuhiro Aoyama, Shogo Yoshihara, Hiroko Takazaki, Takayuki Kato, Hidemasa Goto, Kaoru Mitsuoka, Ryo Nitta, Takao Nakata

## Abstract

Lamellipodia are sheet-like protrusions essential for migration and endocytosis, yet the ultrastructure of the actin cytoskeleton during lamellipodia formation remains underexplored. Here, we combined the optogenetic tool PA-Rac1 with cryo-ET to enable ultrastructural analysis of newly formed lamellipodia. We successfully visualized lamellipodia at various extension stages, representing phases of their formation. In minor extensions, several unbundled actin filaments formed “Minor protrusions” at the leading edge. For moderately extended lamellipodia, cross-linked actin filaments formed small filopodia-like structures, termed “mini filopodia.” In fully extended lamellipodia, filopodia matured at multiple points, and cross-linked actin filaments running nearly parallel to the leading edge increased throughout the lamellipodia. These observations suggest that actin polymerization begins in specific plasma membrane regions, forming mini filopodia that either mature into full filopodia or detach from the leading edge to form parallel filaments. This actin turnover likely drives lamellipodial protrusion, providing new insights into actin dynamics and cell migration.

## Introduction

Lamellipodia, the sheet-like and thin cellular protrusions, are crucial components that form at the leading edge of migrating cells^1,2^. The actin cytoskeleton plays a fundamental role in cell shaping and motility^3^, and which is equally critical in lamellipodia formation and retraction. The lamellipodia-like actin network is a fundamental structure observed in cellular motility, including amoeboid movement and endocytosis^4,5^. These processes are essential for the immune system^6,7^, early development^8^, and cancer metastasis and invasion^9^.

The small GTPase Rac1 localizes the periphery of the plasma membrane and is known as a major upstream factor in lamellipodia formation^10,11^. Rac1 activates the SCAR/WAVE complex^12^, which in turn triggers the activation of the Arp2/3 complex^13^. The Arp2/3 complex, an actin nucleation factor, binds to the sides of existing actin filaments, leading to the growth of new actin filaments at an angle of approximately 70°^14^. This actin branching is believed to be a principal feature of lamellipodia.

Most lamellipodia also possess rib-like structures: microspikes, which do not protrude from the leading edge, and filopodia, which extend beyond it (Fig. 1A). These are straight and bundled actin filaments, reported to be polymerized by the formin/mDia family and crosslinked by fascin^15^.

**Fig. 1.**
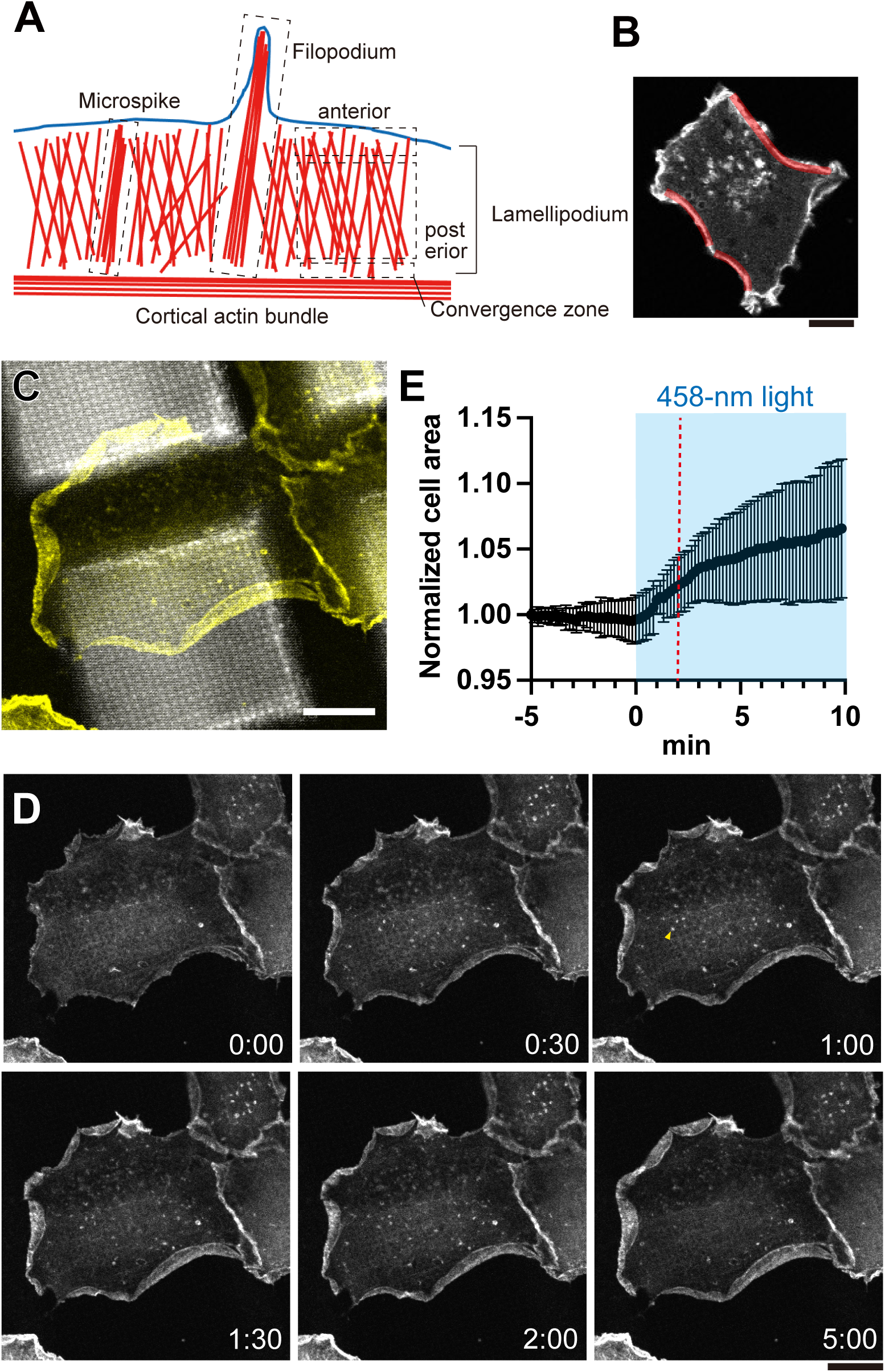
PA-Rac1-induced lamellipodia formation on EM grids. (**A**) Schematic representation of subdomains in lamellipodia. (**B**) Lifeact-mCherry image of COS-7 cells, highlighting the crescent-shaped cell edges in red. Scale bar = 20 µm. **(C–E)** COS-7 cells expressing PA-Rac1 and Lifeact-mCherry on EM grids. Images were captured every 10 sec from −5 min, with PA-Rac1 activated by scanning with a single 458 nm laser within 0–10 min at 10-sec intervals. **(C, D)** Representative time-lapse images. A merged image of DIC and fluorescence of Lifeact-mCherry (yellow) **(c)**, and fluorescence images of Lifeact-mCherry **(D)** are shown. A yellow arrowhead indicates dotted actin structures within the interior of cells Scale bar = 20 µm. **(E)** Quantification of cell area changes. Data are presented as the means ± SD from 6 cells. Blue background indicates the activation period; red dotted line marks the timing of freezing.

The dynamics and the structure of lamellipodia have been extensively studied over the past half-century using both light and electron microscopy (LM and EM). Super-resolution LM, total internal reflection fluorescence microscopy (TIRF), and speckle microscopy have provided insights into protein localization and actin dynamics within lamellipodia^16–21^. However, highly crowded actin structures in lamellipodia make it challenging to resolve single filament structures using LM. In contrast, EM has been instrumental in revealing the ultra-structure of lamellipodia^22–24^. Nonetheless, the chemical fixation and negative staining in conventional EM potentially introduce artifacts. The recent development of cryo-electron tomography (cryo-ET) has overcome these limitations. With advanced image processing, cryo-ET can visualize native structures of proteins and protein complexes in cells^25–27^. Given that lamellipodia are thin cellular structures with a 100–200 nm height, they are ideal targets for cryo-ET. However, observing lamellipodia with intact cell membranes using cryo-ET remains challenging due to sample thickness and ice constraints. Consequently, most cryo-ET studies on lamellipodia have been conducted with fixed and permeabilized cells^28–30^. Moreover, most studies focus on motile cells like fish epidermal keratocytes, observing the structure of already established lamellipodia, which overlooks the early stage of lamellipodia formation. Recently, Chung and colleagues observed lamellipodia with intact cell membranes in spreading cells, utilizing gal-8 for the extracellular matrix (ECM) to facilitate lamellipodia extension thinly enough for cryo-ET analysis^31^. Their methods enable the determination of the polarity of actin filaments and the high-resolution structure of the Arp2/3 complex within an intact cellular membrane. However, protrusions during cell spreading may not fully represent lamellipodia formation.

Here, we employed the optogenetic tool PA-Rac1 (<u>p</u>hoto<u>a</u>ctivatable-Rac1)^32^ to observe lamellipodia formation using cryo-ET. We utilized blue light irradiation within the chamber of an automatic plunge freezer, allowing precise control over the timing of Rac1 activation and its subsequent fixation by quick freezing. The frozen specimens were then subjected to cryogenic correlative light and electron microscopy (cryo-CLEM), and cryo-ET data of lamellipodia with intact cell membranes was obtained. We analyzed the ultrastructure of newly formed lamellipodia with varying degrees of extension, with a focus on the interactions between the plasma membrane and the actin cytoskeleton at the leading edge. These detailed investigations shed light on the ultrastructure of lamellipodia, enhancing our understanding of the mechanisms driving cell motility.

## Results

### Light-induced lamellipodia formation on EM grids

To study the ultrastructure of the actin cytoskeleton and plasma membrane during lamellipodia formation through cryo-ET, it is critical to vitrify cells immediately after stimulation. This ensures that the resultant changes are distinctly observable in subsequent analyses. Accordingly, we utilized the optogenetic tool PA-Rac1 in this study, which consists of the photosensitive LOV2 domain and a constitutively active Rac1 mutant. Blue light exposure changes the LOV2 structure, enabling Rac1 activation, which induces lamellipodia formation^11,32^. For this study, we choose COS-7 cells for their adhesiveness, ease of transfection, and clear observation of lamellipodia formation on EM grids.

Initially, PA-Rac1-induced lamellipodia formation in COS-7 cells was characterized on glass-bottom dishes using confocal laser scanning microscopy (CLSM, Fig. S1A and Video 1). Lifeact-mCherry was used as an F-actin marker^33^. Even before activation by light, cells spontaneously formed lamellipodia (Fig. S1A; 0:00). After activation, more prominent protrusions emerged from all around the cells, clearly differentiating the lamellipodia formation induced by PA-Rac1 (Fig. S1A; 2:30). Significant lamellipodia formation induced by PA-Rac1 was observed along the cell periphery, mainly where the cortex forms inwardly curving arcs, extended over 20 µm in length, and reminiscent of a crescent shape (Fig. 1B). These ‘crescent-shaped edges’ were characterized by abundant cortical actin bundles, marked by the intense fluorescence of Lifeact-mCherry. In line with the hypotheses that lamellipodia formation is initiated by the Arp2/3 complex-mediated side branching from existing actin filaments oriented parallel to the plasmalemma^28^, these regions are reasonable sites of vigorous lamellipodia formation. The initial extension of lamellipodia was typically completed within about 2 minutes of starting blue light irradiation, followed by repeated formation and retraction. The extent of lamellipodia expansion varied among cells and within different areas of the same cell, with the most significant extension reaching lengths of over 5 µm. After the formation of lamellipodia, the actin retrograde flow was observed (Video 1). Microspikes and filopodia, were formed within lamellipodia (Fig. S1A). Interestingly, these structures moved laterally (Fig. S1B), and their tips occasionally detached from the leading edge and gradually collapsed. Eventually, some were incorporated into the pre-existing cortical actin bundles (Fig. S1D). These data suggest that bilateral flow^22^ also occurs in PA-Rac1-induced lamellipodia. Interestingly, the fluorescence intensity of Lifeact-mCherry was notably high at the anterior region of the lamellipodia, comparable to the intensity in cortical actin bundles. In contrast, it significantly decreased towards the central regions within lamellipodia.

Next, lamellipodia formation with PA-Rac1 was induced on EM grids. The formation and mobility of lamellipodia are largely influenced by the ECM. To effectively observe lamellipodia through cryo-ET, they must be thinly spread and close to the substrate. Therefore, EM grids were coated with poly-L-lysine and laminin. Similar to the observations on glass-bottom dishes, lamellipodia formed around the cells upon blue light irradiation, mainly from the crescent-shaped cell edges (Figs. 1C, D and Video 2). The actin retrograde flow and the formation of microspikes and filopodia were also observed, along with their lateral movement (Fig. S1C) and collapsing (Fig. S1E). These observations demonstrate the similarity of lamellipodia formation on EM grids to that observed on glass-bottom dishes, at least through CLSM. Notably, lamellipodia formation from the entire cell perimeter is an essential indicator for identifying PA-Rac1-induced lamellipodia. Another principal indicator is the formation of dotted actin structures within the interior of cells, approximately 1 µm in diameter (Fig. 1D, yellow arrowhead). Although their specific nature remains unidentified. These features allowed us to confidently select the cells with PA-Rac1-induced lamellipodia after plunge freezing.

To determine the optimal timing for freezing, we measured cell area changes post-PA-Rac1 activation (Fig. 1E). The cell area increased significantly within the first four minutes after the start of blue-light irradiation. Despite lamellipodia maintained beyond this time, some areas showed retraction and re-expansion (Video 2). Based on these findings, plunge freezing was performed at the two-minute mark after the start of light irradiation to observe nascent lamellipodia.

### Acquisition of cryo-ET for PA-Rac1-induced lamellipodia

To analyze the ultrastructure of PA-Rac1-induced lamellipodia with an intact membrane, COS-7 cells expressing mVenus-PA-Rac1 and Lifeact-mCherry were cultured on the EM grids and subjected to plunge freezing immediately after two minutes of blue light irradiation (Fig. 2). Cells with PA-Rac1-induced lamellipodia were selected based on the previously mentioned criteria through Leica Cryo-CLEM light microscopy. Subsequently, these grids were subjected to cryo-EM, and tilt series were collected while correlated with the cryo-LM images (Fig. S2). Since the plasma membrane tended to deform at the holes of the Quantifoil (Figs. S3A, B)^34,35^, these areas were avoided for analysis. Although the signal-to-noise (S/N) ratio was reduced on the carbon sheet, there was still sufficient contrast to segment cellular structures (Figs. S3C, D). After several attempts of grid preparation and data collection, we successfully prepare well vitrified two grids, and measured by cryo-EM. We succeeded in analyzing 16 tomograms from three cells, capturing images from the leading edge to the rear regions of the lamellipodia with varying degrees of extension (Figs. S2, S4). We measured the variety of potential lamellipodia regions indicated by white squares in Fig. S4 and analyzed them.

**Fig. 2.**
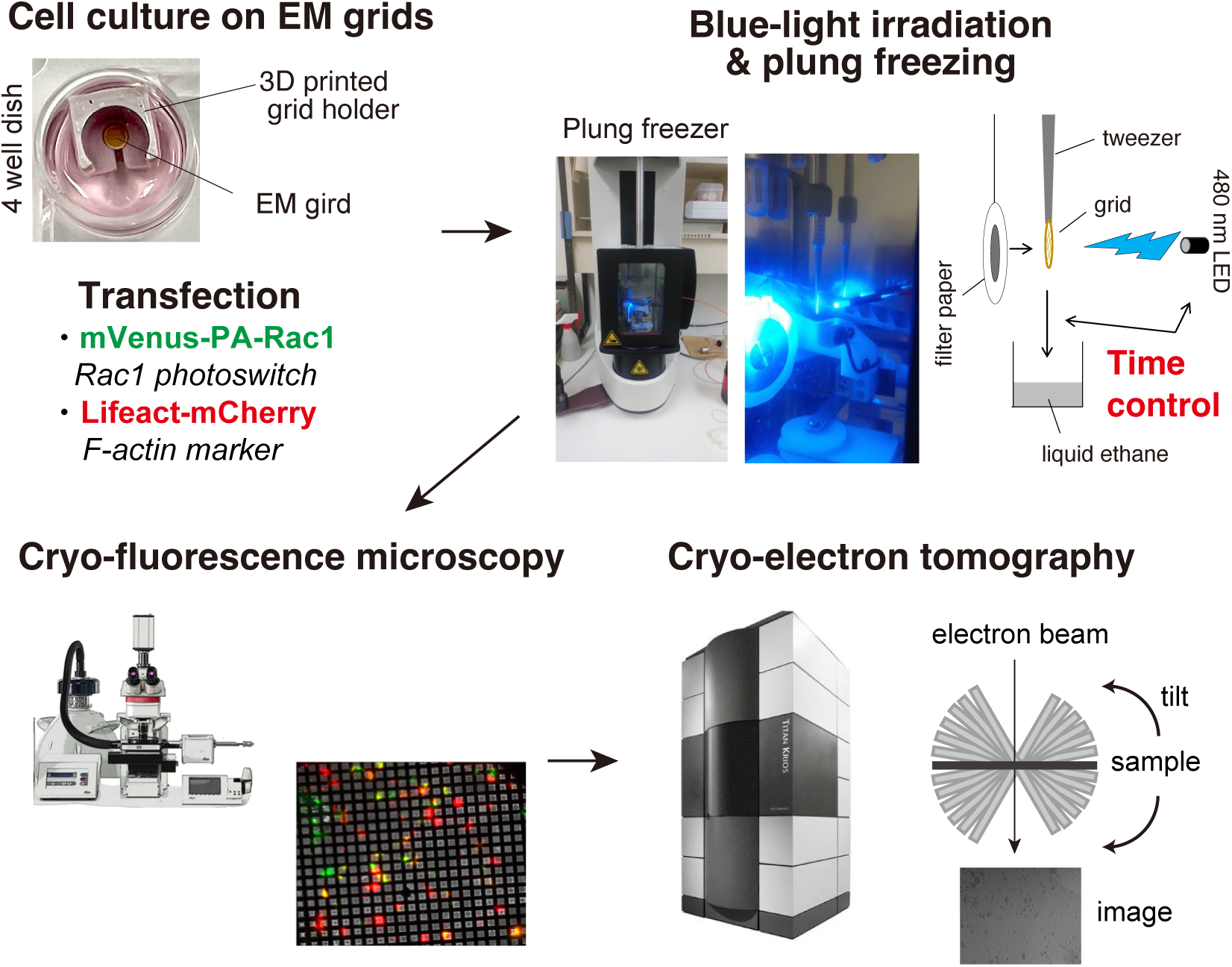
Experimental workflow from sample preparation to tomogram acquisition. Laminin-coated EM grids were placed on the 3D-printed grid holders in 4 well dishes, and cells were seeded on the EM grids. At the same time, plasmids encoded mVenus-PA-Rac1 and Lifeact-mCherry were transfected into the cells. The next day, the cells were rapidly frozen in a plunge freezer after inducing lamellipodia formation by activating PA-Rac1 with blue light irradiation, controlling the timing of the freezing. The frozen cells were observed using cryo-fluorescence microscopy to identify the cells that had formed lamellipodia. By correlating the fluorescence images with low-magnification electron microscopy images, the regions for tomography were determined. Subsequently, continuous tilt series images were acquired by cryo-electron microscopy from −70 degrees to +70 degrees, and tomography was reconstructed.

The average height of lamellipodia from three cells ranged from 50 to 250 nm (Fig. S5A), and filament density was between 2,500 and 15,000 per um^-3^ (Fig. S5B), aligning with previous studies^31,36^. The lower actin filament density in Cell 2 might be related to pronounced lamellipodia spreading. Across all lamellipodia, filaments perpendicular to the leading edge were relatively rare, peaking at angles of around 20–30 degrees and around 90 degrees to the leading edge (Fig. S5C). When analyzed by individual cells, the orientation of actin filaments varied from cell to cell (Fig. S5D). The orientation of actin filaments, including regions outside the lamellipodia, was depicted in Fig. S5E. We also explored the correlation between the length of actin filaments and their orientation to the leading edge (Fig. S5F). For filaments shorter than 200 nm, a peak orientation was 15 to 25 degrees. In contrast, longer filaments tend to align more parallel to the leading edge. The Arp2/3 complex mediates actin branch formation, crucial for organizing the actin cytoskeleton in lamellipodia^20,23,30^. We identified actin branches by meticulously reviewing tomograms within the 3dmod software, using the segmented actin filaments as references. The criteria for identifying branches were based on their position at the ends of filaments, branching angle around 70 degrees, and detecting densities on the base of branches, indicative of the Arp2/3 complex. Identified actin branches are represented as black dots in the segmentation images, with representative examples displayed in Fig. S6A. The average frequency of the actin branch was one branch per 6.2 µm of filament length (Fig. S6B). On the other hand, the average frequency of the actin branch outside of lamellipodia was one branch per 31.7 µm filament length, indicating approximately five times more branches within the lamellipodia. These data underscore the essential role of actin branches in lamellipodia formation.

### The architecture of the actin cytoskeleton and protrusive membrane structure of the growing lamellipodia

Next, we analyzed the details of the actin cytoskeleton at various stages of lamellipodia formation, with the assumption that different degrees of extension represent different stages of lamellipodia formation. First, we examined a lamellipodium with a relatively low degree of extension, measuring approximately 0.5 µm (Fig. 3). The actin bundle on the left side in the tomography (Figs. 3D, E and Movie 3) could be the original cortical actin. Another example of such a degree of extended lamellipodia is shown in Fig. S4C-1. Focusing on the leading edge, the plasma membrane appeared slightly undulated. Here, we classified the surface features into three groups based on the ratio of height to width (H/W) of the protrusions (Fig. S7). To determine H and W, we manually selected all visibly distinguishable protrusions with a height of 20 nm and above. The width (W) was measured as the distance between the two endpoints at the base of each protrusion, while the height (H) was defined as the perpendicular distance from this base line to the apex of the protrusion. Based on these measurements, we categorized the protrusions as “Large protrusion” (H/W > 0.5), “Minor protrusions” (0 < H/W ≤ 0.5). The remaining areas, where H/W < 0, were classified as “Concave regions” (Fig. S7A). At the Minor protrusions, several actin filaments ran towards the leading edge (Figs. 3F, G). Most of these actin filaments, located just beneath the protrusive plasma membrane, had their growing ends oriented toward the leading edge. In these Minor protrusions, the actin cytoskeleton was not well-organized, lacking cross-linked actin filaments, and slightly curved filaments were observed. Inside the lamellipodium, actin filaments were oriented in various directions, showing no distinctive structures.

**Fig. 3.**
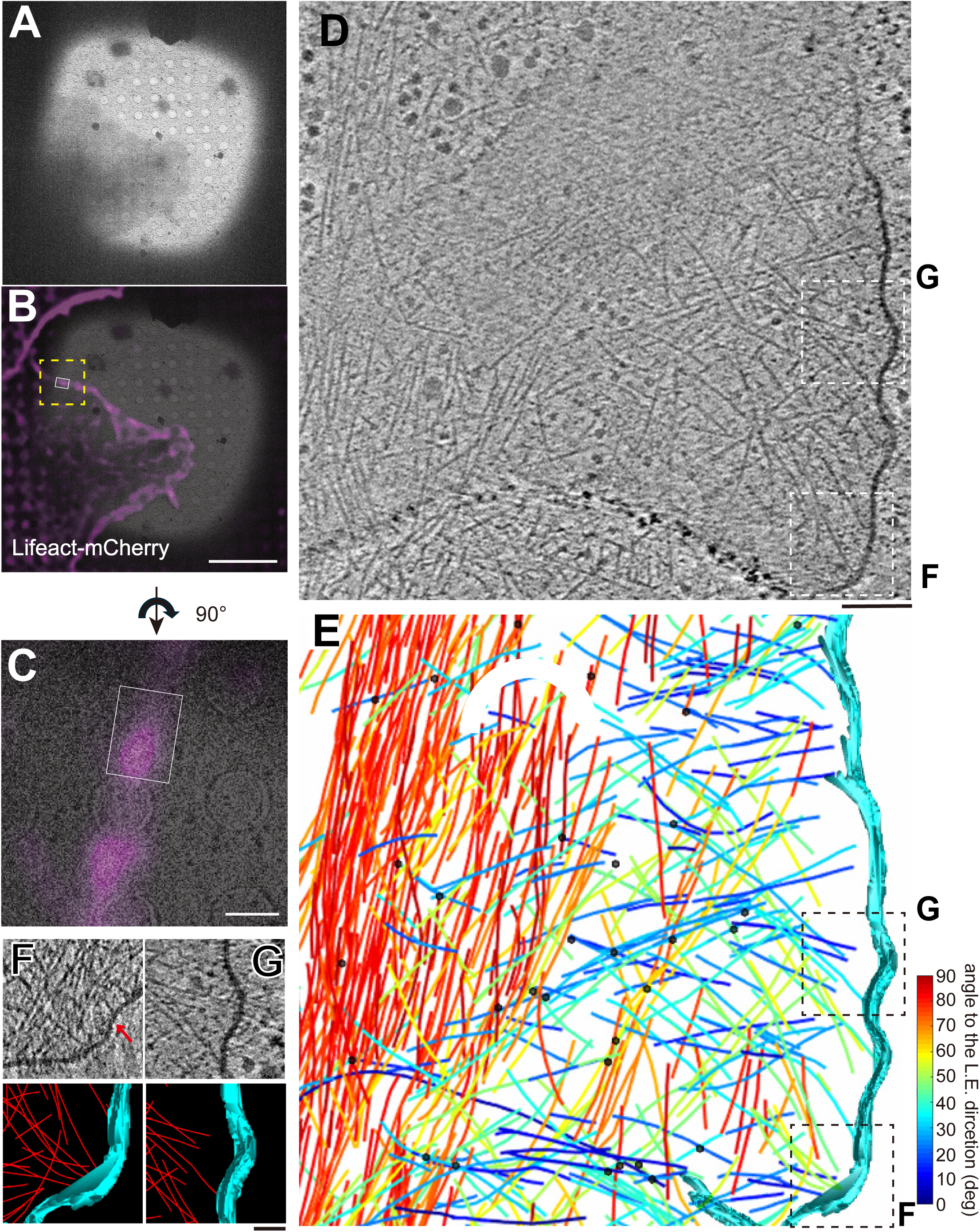
Cryo-ET of PA-Rac1-induced lamellipodia in the early stage of formation. **(A–C)** A cryo-EM image **(a)** and corresponding cryo-CLEM images **(B, C)** of a PA-Rac1-induced lamellipodium in Cell 1 (Fig. S2) extended approximately 0.5 µm. (**C)** An enlargement of the area within the yellow rectangles in **b**. Scale bar = 10 µm in **b** and 2 µm in **C**, respectively. **(D, E)** A cross-sectional (x-y) slice from cryo-ET **(D)** and a corresponding segmented image **(E)** obtained from the area marked by a white box in **B** and **C**. The plasma membrane is represented in cyan. Actin filaments are color-coded to indicate their orientation relative to the leading edge (L.E.) of the cell. Here, 0 degrees represents perpendicular orientation relative to the leading edge, while 90 degrees indicates parallel orientation. Branching points of actin filaments are marked with black dots. Scale bar = 200 nm. **(F, G)** Enlarged images of the protrusive structure at the leading edge shown in the black dotted box in **D** and **E**. Scale bar = 50 nm.

Next, we studied a lamellipodium extended to approximately 1.5 µm (Fig. 4 and Movie 4). In this lamellipodium, two Large protrusions were observed at the leading edge (Figs. 4F, G). In the Large protrusion highlighted in Fig. 4F, the growing end of actin filaments localized in a region of the plasma membrane. These filaments formed two clusters, with some filaments running parallel and extending in different directions, guiding the membrane protrusion along their paths. In contrast, in the Large protrusion observed in Fig. 4G, the actin cytoskeleton was arranged in a manner that two or more straight actin filaments ran parallel and oriented toward the leading edge, similar to that in microspikes and filopodia but on a smaller scale. The spacing between parallel actin filaments in these protrusions (Fig. 4H), approximately 8 nm, suggested fascin-mediated cross-linking, characteristic of microspike and filopodia formation^37,38^. Due to the limited number of filaments and their shorter length (Fig 4E), hereafter, we refer to this structure as “mini filopodia.” Such structures were also detected in another tomogram (Fig. S4C-2).

**Fig. 4.**
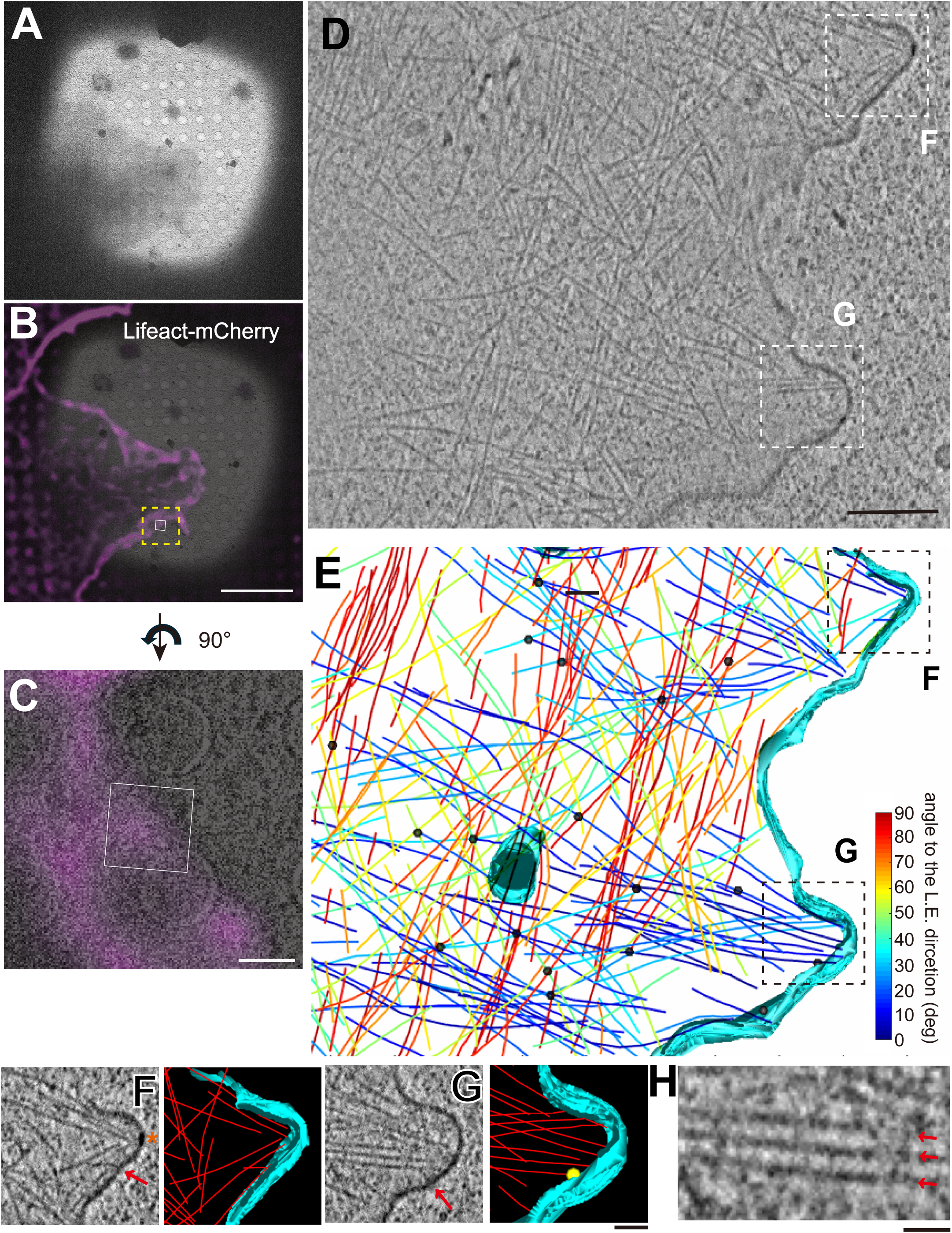
Cryo-ET of PA-Rac1-induced lamellipodia in the middle stage of formation. **(A–C)** A cryo-EM image **(A)** and corresponding cryo-CLEM images **(B, C)** of a PA-Rac1-induced lamellipodium in Cell 1 (Fig. S2) extended approximately 1.5 µm. **(C)** An enlargement of the area within the yellow rectangles in **B**. Scale bar = 10 µm in **B** and 2 µm in **C**, respectively. **(D, E)** A cross-sectional (x-y) slice from cryo-ET **(D)** and a corresponding segmented image **(E)** obtained from the area marked by a white box in **B** and **c**. Scale bar = 200 nm. **(F, G)** Enlarged images of protrusive structure at the leading edge shown in the black dotted box in **D** and **E**. Yellow dots mark branched points of actin filaments; red arrows point locations where actin filaments attach to the plasma membrane; an orange asterisk denote regions where the plasma membrane shows high intensity. Scale bar = 50 nm. **(H)** Enlarged images of the actin filaments running parallel in **G**, with three filaments indicated by red arrows running parallel to each other. Scale bar = 10 nm.

Notably, within these Large protrusions, the localized high density areas on the plasma membrane were observed (Fig. 4F, orange asterisk). These areas suggest the presence of protein complexes or specialized membranes, offering a promising direction for further exploration. Meanwhile, in the Minor protrusion of the leading edge (below box f in Fig. 4E), several actin filaments ran towards the leading edge without bundling, like the minor protrusions in Fig. 3. In contrast, some single filaments ran towards the leading edge in the Concave region, with more prominent filaments running parallel to the leading edge in these regions.

When focusing on the interior of the lamellipodium (Fig. 4E), long actin filaments parallel to the leading edge were prominent. Additionally, a few actin filaments color-coded in blue at the lower left of the tomogram appeared detached from the leading edge but exhibited an orderly structure like mini filopodia. For other actin filaments, the similar orientation of neighboring filaments suggests forming an organized actin network in this degree of extended lamellipodia.

Next, we quantified the distance between the growing ends of actin filaments and the plasma membrane for each of the three groups, explicitly focusing on actin filaments oriented toward the leading edge (Figs. S7B–E). Actin filaments adhering to the plasma membrane (red arrows in Figs.4F, G, and Figs. S7C, D) were relatively common in the Concave regions, accounting for about half of the observed cases. On the other hand, in both Large and Minor protrusions, these adherent filaments constituted only about 15% of the total, with a notable number of filaments showing gaps between their growing ends and the plasma membrane. These gaps predominantly ranged from 10 to 20 nm, although no uniform distance was consistently observed.

### Analysis of actin architecture in extended lamellipodia

In our tomograms, the lamellipodia in Cell 2 was most extended, measuring over 3 µm (Fig. 5). In the LM image, a few filopodia or microspikes can be observed within the lamellipodia (Figs. 5B, C). Part of filopodium was detected in a tomogram (Figs. 5D, E and Movie 5), where bundles of over ten filaments ran linearly toward the leading edge. When magnified, these bundles were spaced approximately 8 nm apart (Fig. 5H). Other actin bundles ran parallel to the leading edge inner lamellipodia (for example, white/black dotted boxes in Figs. 5F, G and Movie 6). These three to five actin filaments were also cross-linked at approximately 8 nm (Figs. 5I, J). These data suggest that these actin bundles, including filopodia and mini filopodia, are formed by the exact mechanism involving fascin-mediated cross-linking.

**Fig. 5.**
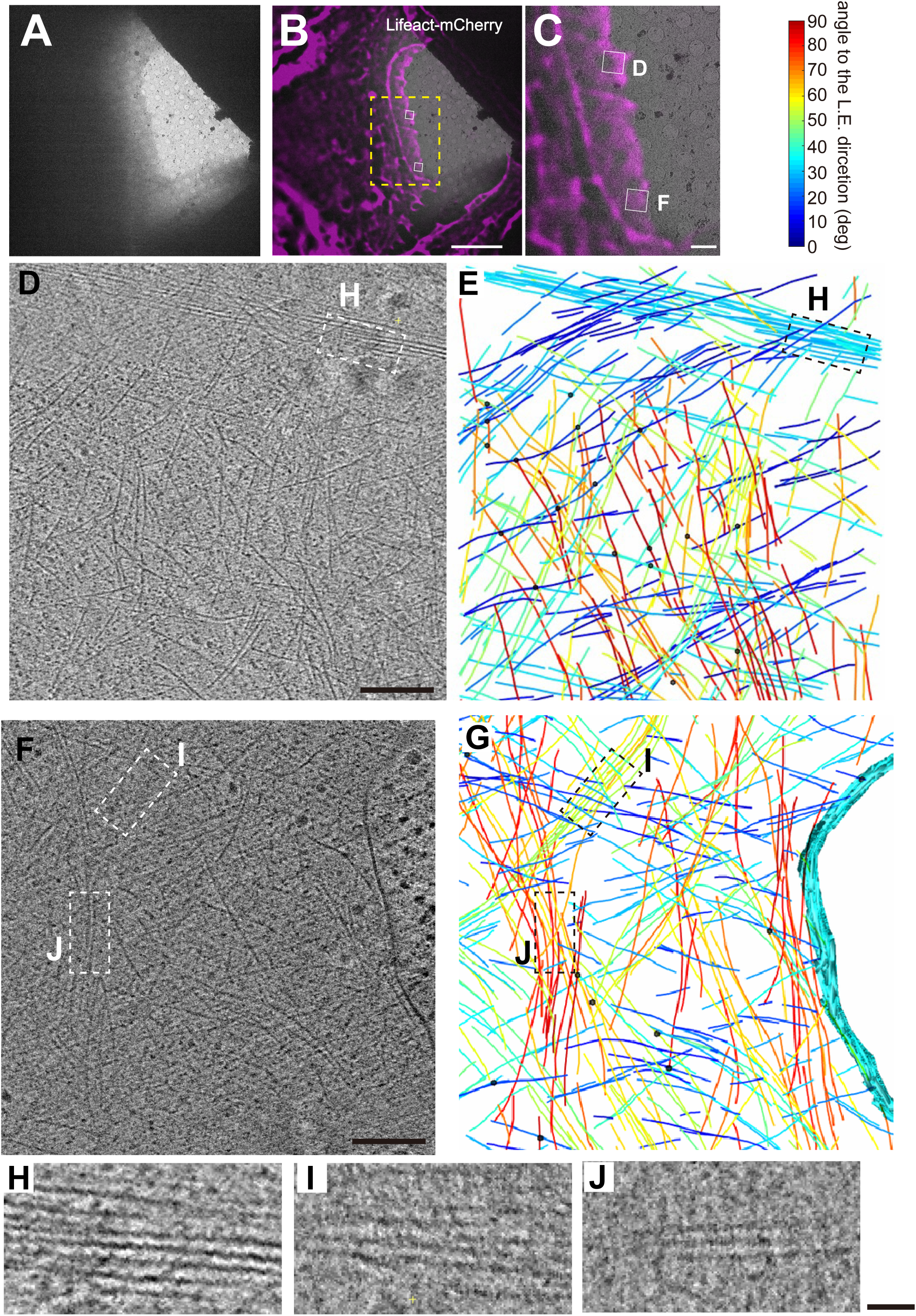
Cryo-ET of PA-Rac1-induced lamellipodia in the late stage of formation. **(A–C)** A cryo-EM image **(A)** and corresponding cryo-CLEM images **(B, C)** of a PA-Rac1-induced lamellipodium in Cell 2 (Fig. S2) extended over 3 µm. **(C)** An enlargement of the area within the yellow rectangles in **B**. Scale bar = 10 µm in **B** and 2 µm in **C**, respectively. **(D–G)** Cross-sectional (x-y) slices from cryo-ET **(D, F)** and corresponding segmented images **(E, G)** obtained from the area marked by white boxes in **B** and **C**. Scale bar = 200 nm. **(H–J)** Enlarged images of actin bundles in filopodia (dotted boxes in **D** and **E**), and actin bundles running parallel to the L.E. within lamellipodia (dotted boxes in **F** and **G**). Scale bar = 10 nm.

Our observations primarily highlight the role of fascin in the prominent actin cross-linking within lamellipodia. Detailed analysis revealed variations in actin filament density, with regions of densely packed filaments and areas where they were sparse (Fig. S4). Many closely situated actin filaments ran in similar directions, hinting at the involvement of other actin cross-linking proteins, such as alpha-actinin^39^, in their organization.

The protrusive membrane at the leading edge was not captured in this cell. In the Concave regions of the leading edge, a single filament ran toward it (Figs. 5F, G), similar to previous observations. Additionally, prominent actin filaments ran parallel to the leading edge beneath the plasma membrane in these regions.

### Comparative analysis of actin dynamics and structure in anterior and posterior regions of lamellipodia

Here, we obtained tomograms in various regions from the anterior to the posterior regions of lamellipodia. The anterior region of the lamellipodia, located about 1 to 1.5 µm from the leading edge, is a specialized area where the activity of actin polymerization and depolymerization are exceptionally high^1^. Moreover, proteins that regulate actin dynamics, such as Ena/VASP, the SCAR/WAVE complex, and profilin, concentrate in the anterior region. Therefore, the anterior region is frequently analyzed separately from the more posterior parts of the lamellipodia (Fig. 1A). In Cell 3, the high fluorescence intensity of Lifeact-mCherry at the anterior regions and significantly lower intensity in the posterior regions of lamellipodia (Figs. 5B, C), suggesting that there could be structure differences between these regions. To compare these differences, we classified tomograms, including other cells, based on distance from the leading edge: ‘anterior’ (near) and ‘posterior’ (far) (Fig. S8A). In the anterior region, the orientation of actin filaments was similar to the overall one (Fig. S8B). On the other hand, in the posterior region, the peak at 30 to 40 degrees was more pronounced, with another peak observed around 90 degrees. Although the density of actin filaments appeared to decrease towards the rear of lamellipodia (Fig. S8C), this reduction was surprisingly minor compared to the variance observed in Lifeact-mCherry fluorescence. Both regions contained fascin-bundled actin filaments. Despite differences in orientations, no obvious distinction in the actin cytoskeleton between these regions was identified, except for structures associated with the plasma membrane at the leading edge.

### The ultrastructure of the convergence zone of lamellipodia and cortical region of cells

The convergence zone of the lamellipodia, located near the original cortical region of the cells^40,41^, exhibited a predominantly unorganized actin cytoskeleton with minimal branching and a high degree of filament bendiness (Fig. 6 and S4 B6–B8). The cortical actin bundles showed continuous filaments extending across entire tomograms, indicating their composition of actin filaments exceeding 1 µm in length (Fig. 6B). Within these bundles, dozens of actin filaments twisted together with few branching. Additionally, microtubules running parallel to the actin bundles suggested a coordinated structural arrangement.

**Fig. 6.**
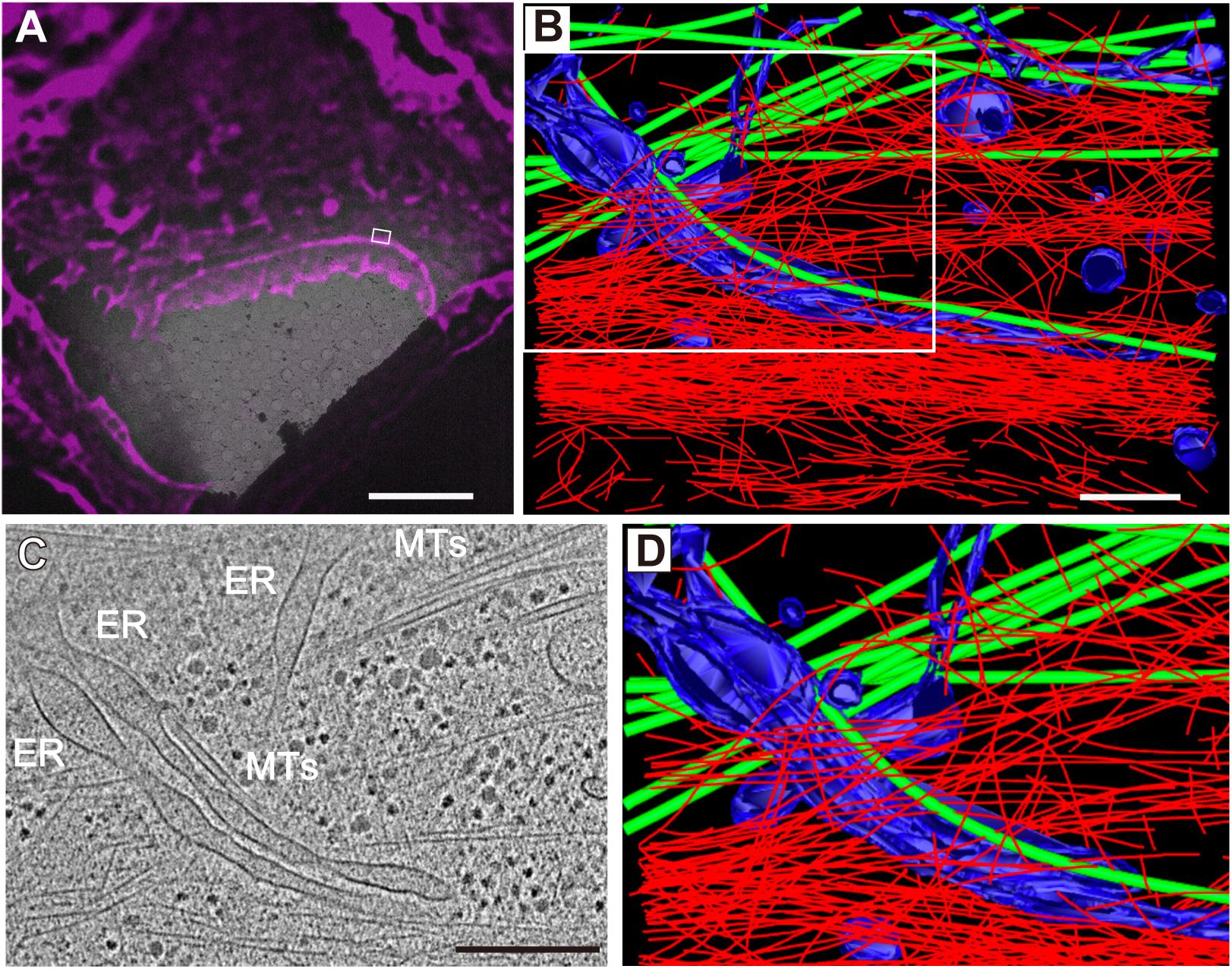
Cryo-ET of the rare region of lamellipodia and cell cortical region before lamellipodia formation. **(A)** Cryo-CLEM image of the same cell shown in Fig. 2, cryo-ET area in white box. Scale bar = 10 µm. (**B**) Segmented cryo-ET of white box area in **A**. Blue: ER and inner cellular membrane structure, Green: Microtubules, Red: actin filaments. Scale bar = 200 nm. (**C, D**) Magnified view of the area in B, focusing on the parallel arrangement of tubular ER and microtubules. Includes a cryo-ET slice **(C)** and its segmentation **(D)**. Scale bar = 200 nm.

Beyond these bundled actin filaments, tubular membrane structures were observed alongside microtubules, indicative of the endoplasmic reticulum (ER; Figs. 6C, D). This arrangement aligns with previous studies in the LM^42^, highlighting the role of microtubules and ER in cellular transport and the maintenance of structural integrity.

## Discussion

In the present study, we conducted ultrastructural analyses on lamellipodia formation in cells with intact plasma membranes, using a combination of cryo-ET and optogenetics. The density and the orientation of actin filaments (Fig. S5A, B) were consistent with previous studies^22,31,36^, affirming our methodology. Furthermore, the preservation of protrusive membrane structures and internal ER membrane (Fig. 6) suggests the effectiveness of the freezing process.

One of the challenges in this study was the limited number of cells suitable for cryo-ET observation. The extended blotting process required for cryo-ET preparation can desiccate cells, risking cell death. Indeed, we had to discard nearly half of the preparation grids because the cells were deemed dead upon examination with cryo-LM. Moreover, even among surviving cells, many exhibited blebbing–signs of stress or damage. Consequently, when selecting grids containing healthy cells, only those with adequately thin ice layers for practical cryo-EM observation were chosen, comprising less than 20% of all prepared grids. This limitation resulted in a smaller sample size for analysis.

Despite this, the precise control of lamellipodia formation using optogenetics allowed us to uncover novel ultrastructural details that differ from previous observations in motile keratocytes^28,29,43^ or during spreading cells post-plating^31^. These findings raises fundamental questions about the organization and role of the actin cytoskeleton during lamellipodia formation. The frequency of actin branches (Fig. S6B) was significantly lower, about one-tenth compared to previous studies^28,31,43^. While it is conceivable that some branches were not detected due to the inherently noisy images produced by cryo-ET, it is unlikely that the number of missed branches would account for a tenfold difference. This discrepancy could be attributed to distinct experimental conditions, including the type of cells and protrusive activity of lamellipodia. The density of actin branching mediated by Arp2/3 is believed to influence the stiffness of the actin cytoskeleton and protrusion speed of lamellipodia^44,45^. Even considering these discrepancies, the higher frequency of actin branches found within the lamellipodia affirms their critical roles in the organization of the actin cytoskeleton during lamellipodia formation.

Taking advantage of preserving intact cell membranes, we analyzed the architecture of the plasma membrane and the actin cytoskeleton at the leading edge in detail. Based on our observations, we propose a model for the reorganization of the actin cytoskeleton during lamellipodia formation, as illustrated in Fig. 7, encompassing different stages of extension.

**Fig. 7.**
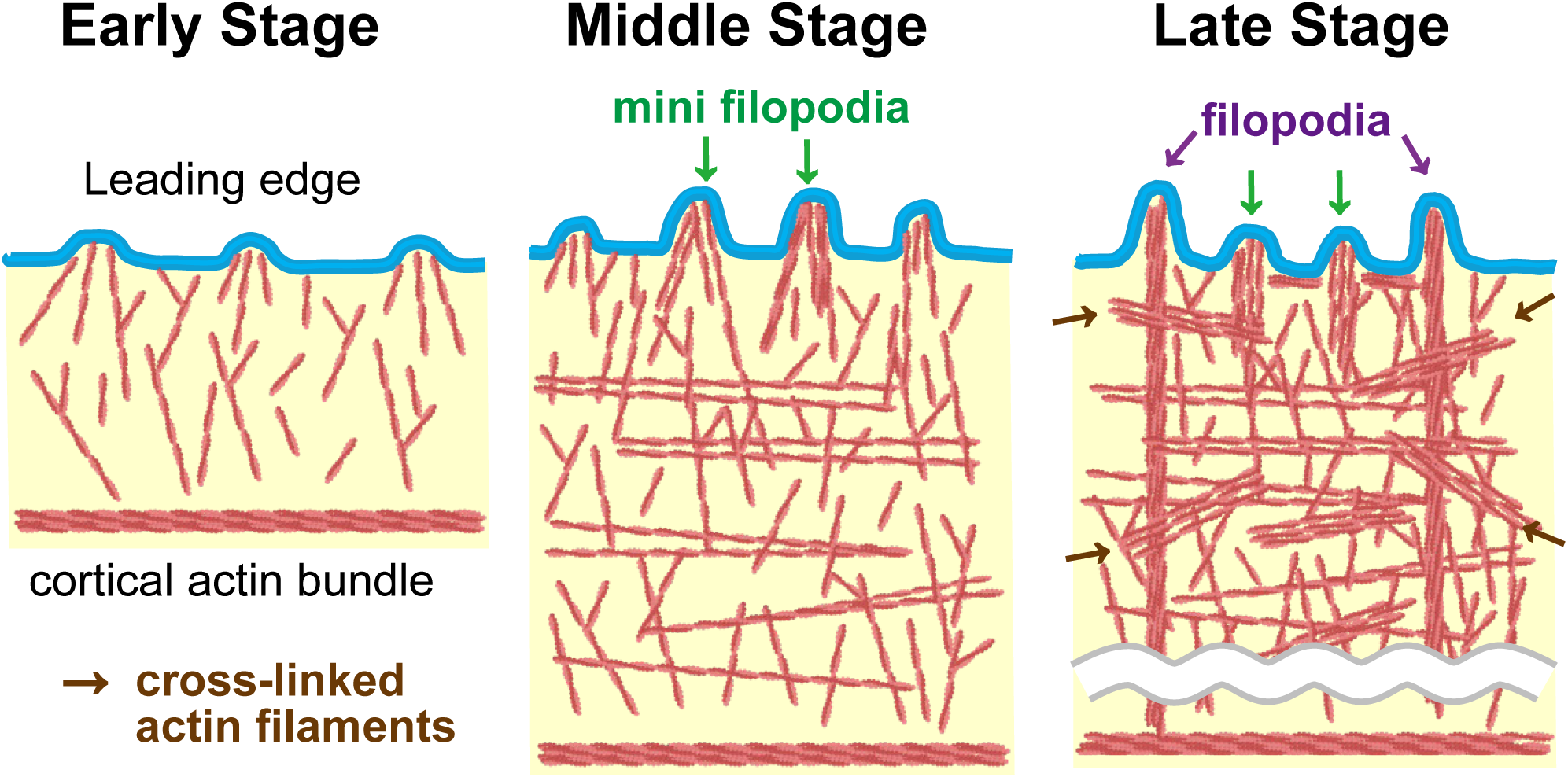
A model of actin cytoskeleton reorganization during formation of lamellipodia. This figure illustrates a model for the reorganization of the actin cytoskeleton during lamellipodia formation, comprising three stages: **Early Stage**: Minor protrusions form, with unbundled actin filaments growing towards the leading edge, **Middle Stage**: mini filopodia emerge, with some actin filaments becoming cross-linked. **Late Stage**: matured filopodia are present, with cross-linked actin filaments running nearly parallel to the leading edge throughout the lamellipodia. See Discussion for details. Illustration created with BioRender.

In the early stage, Minor protrusions form at the leading edge (Fig. 3 and Movie 3). Several actin filaments grew towards the leading edge but were not bundled (Figs. 3F, G). Conversely, in Convaved regions, actin filaments grow towards the leading edge individually. The localized actin polymerization in the specific regions suggests the involvement of liquid-liquid phase separation (LLPS) of actin-polymerizing factors^46–48^. Meanwhile, no organized actin structures were observed within the inner regions of lamellipodia at this stage.

In the middle stage, the actin cytoskeleton exhibited a more structured and organized arrangement compared to the early stage (Fig. 4 and Movie 4). Large protrusions appeared, with 5 to 10 actin filaments growing to the leading edge, some of which were cross-linked at approximately 8 nm intervals. These structures, which we refer to as mini filopodia, resemble microspikes and filopodia, but have fewer and shorter filaments, making them detectable only with EM. The formation of microspikes and filopodia within lamellipodia is known to be facilitated by Ena/VASP proteins^49^. The spacing between actin filaments in mini filopodia suggests bundling by fascin, which enhances Ena/VASP activity^50^, indicating a cooperative mechanism in the protrusion formation. Studies on LLPS have shown that actin filaments become bundled after nucleation^39,48^, suggesting that mini filopodia form through the growth and bundling of Minor protrusions. At this stage, Minor protrusions continue to be observed, suggesting that these structures turn over during lamellipodia formation. Additionally, there was a mini filopodia-like actin structure near the leading edge that appeared to be separated from it, suggesting it could have detached after formation. Furthermore, some long actin filaments ran nearly parallel to the leading edge within the inner regions of lamellipodia.

In the late stage, tomograms did not capture protrusive structures at the leading edge in this study. Instead, prominent microspikes and filopodia were observed in cryo-LM, and parts of these structures were captured in a tomogram (Fig. 5). These microspikes and filopodia are considered to form as a result of further maturation of mini filopodia. Conversely, many bundled actin filaments running nearly parallel to the leading edge were observed within the inner regions of lamellipodia (Fig. 5I, J). It is possible that these bundled actin filaments have originally been part of mini filopodia and later detached from the leading edge. In living cells, microspikes and filopodia moved laterally due to bilateral flow, occasionally collapsing (Figs. S1B-E). Such phenomena could also occur in mini filopodia, suggesting that mini filopodia might either mature into filopodia or detach from the cell membrane to form filaments running nearly parallel to the leading edge. Actin filaments running parallel to the leading edge were often longer (Fig. S5F). This makes sense as the retrograde flow of actin tends to push longer filaments into obstacles where they stay. This is akin to how longer logs in a river are more likely to encounter obstacles and align parallel to the flow. Within cells, these obstacles might be other actin filaments and premature cellular adhesions^51^.

The mechanism of lamellipodia formation is as follows: the initiation of actin polymerization at the specific region of the plasma membrane at the leading edge leads to the formation of Minor protrusions. These Minor protrusions then develop into mini filopodia through the growth and bundling of actin filaments, which drive the plasma membrane forward. These structures can further evolve into filopodia, but sometimes detach from the leading edge and remain within the lamellipodia as actin bundles parallel to the leading edge. The turnover of these structures within the lamellipodia is a major driving force for lamellipodia extension. Our model complements the traditional role of Arp2/3-mediated branching in lamellipodia extension.

Both in Large and Minor protrusions, gaps existed between the growing ends of actin filaments and the plasma membrane (Fig. S7E). These gaps are expected, as actin polymerization requires free space for the attachment of actin monomers to the filament tips. The emergence of these spaces can be attributed to the random movement of the plasma membrane, known as Brownian motion^52^. At the growing plus ends of actin filaments, proteins like Ena/VASP and mDia/formin are believed to function, but the resolution of our cryo-ET was insufficient to identify any complex densities. Conversely, in the Concave regions, more actin filaments directly contacted the plasma membrane than protrusion regions (Figs. S7D, E), likely due to higher tension in these areas. Moreover, some actin filaments ran parallel to the leading edge beneath the plasma membrane, likely helping to resist membrane tension.

Cryo-LM played a crucial role in identifying cells with PA-Rac1 induced-lamellipodia and documenting their states. In particular, Cell 2, which exhibited the most extended lamellipodia, provided significant information. Tomography No.1 and No.2 captured the anterior region (Figs. S4B-1,2), with high Lifeact-mCherry fluorescence. In contrast, tomography No.3 to 5 captured the posterior region (Figs. S4B-3–5), showing diminished fluorescence. Despite differences in fluorescence intensity, the density of actin filaments was not as varied (Fig. S5B). This discrepancy could be due to two factors. First, at the tips of the lamellipodia, actin polymerization and depolymerization are active, with abundant G-actin^16,53^. Lifeact-mCherry is known to bind not only to F-actin but also to G-actin^33^. Second, actin retrograde flow could carry Lifeact-mCherry away from the posterior region^54^. Although actin filament orientation differed between regions (Figs. S8A, B), the limited number of tomograms prevented clear elucidation of specific structural differences.

In this study, we used the optogenetic tool PA-Rac1 to induce lamellipodia formation. Optogenetics allows for precisely manipulating intracellular signaling molecules with light, providing high specificity and time control^55,56^. This method addresses a fundamental limitation of electron microscopy: the inability to observe temporal dynamics due to specimen fixation. Although we froze samples at a single time point, the combination of light irradiation and rapid freezing allows for precise control over reaction times (Fig. 2), creating pseudo-time-lapse cryo-samples. By examining these frozen specimens, we will be able to investigate the dynamics of lamellipodia formation at nanometer resolution. Previous studies have employed optogenetics with cryo-EM to explore dynamic changes in neural cells^57,58^ and purified proteins^59^, demonstrating the potential of integrating temporal control with structural analysis. However, our study uniquely applies this approach to in-cell cryo-ET without staining, enabling detailed observation of the ultrastructure in intact, living cells during lamellipodia formation. Future studies can build on this approach by increasing the number of biological replicates and varying the duration of stimulation to confirm the sequence of events proposed here. The ability to use optogenetics for time-resolved structural analysis holds promise for further elucidating the dynamic processes of actin reorganization in lamellipodia, providing a powerful tool for future investigations.

## STAR★Methods

### Cell culture

COS-7 cells (RCB0539: RIKEN BRC through the National BioResource Project of the MEXT/AMED, Japan) were cultured in Dulbecco’s modified Eagle’s medium (DMEM, Nacalai Tesque #08458-65), supplemented with 10 % fetal bovine serum (FBS, Biowest, #S1810) and incubated at 37°C with 5% CO_2_.

Before seeding the cells, 200 mesh gold holey carbon grids (R1.2/1.3; Quantifoil Micro Tools) were coated with 0.01% (w/v) poly-L-lysine (Nacalai Tesque, #28360-14), applied with 20 nm gold colloidal markers (Sigma-Aldrich, #741965), and coated with thin carbon as described previously^35^. The grids were again coated with 0.01% (w/v) poly-L-lysine for over 1 hour and coated with 5 µg/mL laminin (Thermo Fisher Scientific, #23017015). A total of 1×10^5^ COS-7 cells were seeded onto the grids, which were placed in 3D-printed grid holders^60^ in a four-well plate (SPL Life Sciences, #30004).

The transfection mixture, composed of 1 µg of pTriEx-mVenus-PA-Rac1^32^ (kindly gifted from Klaus Hahn; Addgene plasmid, #22007) and 0.02 µg of Lifeact-mCherry^61^, 50 µl of Opti-MEM^TM^ (Thermo Fisher Scientific #31985070), and 3.6 µl of ViaFect^TM^ transfection reagent (Promega, #E4982), was incubated at RT for 15 minutes. This transfection mix was added to the cells just before seeding.

After transfection, the cells were incubated at 37°C with 5% CO_2_ for 16–20 hours before vitrification. Before vitrification, the expression of the fluorescent proteins was confirmed using either a BZ-X700 fluorescence microscope (Keyence) or an EVOS M5000 imaging system (Thermo Fisher Scientific).

For observing live cell imaging on the glass-bottom dishes, cells were cultured and transfected according to previously described methods ^62^. Briefly, cells were seeded on glass-bottomed dishes (Greiner Bio-one, #627870) coated with collagen (Cellmatrix Type IC; Nitta Gelatin, #631-00771) two days before the observations. The following day, cells were transfected with the plasmid described above, and analyses were conducted a day later.

### Live cell imaging

The grids with cultured cells were inverted so that the cell side faced downward and were then submerged in a glass-bottom dish (Violamo, #GBCD15) filled with Leibovitz’s L-15 (Thermo Fisher Scientific, #11415064). Cells were analyzed under the serum-starved condition in a sealed, heated chamber during live cell imaging at 37 °C without CO_2_ supplementation. Images were obtained using a confocal laser scanning microscope (FV1200, Olympus) on an IX83 microscope (Olympus) equipped with 40×/0.95 NA dry objective lenses and FV10-ASW software (Olympus). Photoactivation and imaging utilized standard setting for ECFP(C-Y-R), EYFP(C-Y-R), DsRed2(C-Y-R), and DIC (Argon laser power: 5% at 458-nm and 1 % at 515-nm; diode laser power: 5% at 559-nm; DM 458/515/560 dichroic excitation, SDM510 and SDM560 emission filters; 475–500 nm emission window for ECFP, 530–540 nm emission window for EYFP, and 570–670 nm emission window for DsRed2; DIC images were obtained with DsRed2 settings) using the sequential line capturing mode (scan rate: 10 µs per pixel, pixel size 4.83 µm). The interval for both imaging and optoactivation was set to 10 seconds. PA-Rac1 activation was induced using the ECFP channels. mCherry images were acquired through the DsRed2 channel. Separately, mVenus images were captured with the EYFP channels only after live imaging to confirm the expression of mVenus-PA-Rac1.

Image analysis was conducted using ImageJ/Fiji software (NIH)^63^. The changes in cell area were quantified using mCherry images. These images were binarized and cell area was measured using the “Analyze Particles” function. Adobe Photoshop 2024 and Illustrator 2024 (Adobe Systems) were used for the final figure preparation.

### Blue light irradiation and vitrification

Samples were vitrified using either a Leica GP2 plunger (Leica Microsystems) or the Vitrobot Mark IV System (Thermo Fisher Scientific) set to 37°C and 95% humidity. Before transfer into the blotting chamber, the medium was manually blotted off, and 5 µl of PBS was added to the grids. Inside the blotting chamber, blue light irradiation was performed using a fiber-coupled LED (470 nm, 1000 mA; Thorlabs, #470F1), which was connected to a ferrule patch cable (φ400 µm core, 0.39 NA; Thorlabs, #M79L01). The output of LED was regulated by a T-Cube LED driver (1200 mA; Thorlabs, #LEDD1B). The cable was fed into the blotting chamber through an aperture reserved for sample application, with the blue light projecting directly from the front of the grid at approximately 1 cm distance, at maximum power. The irradiation timing was manually managed to ensure that blotting was completed, and the sample was plunged into liquid ethane precisely 2 minutes after the onset of irradiation. Vitrification of the grids in liquid ethane (−185°C) was following backside blotting for 10 seconds or bilateral blotting for 5 sec, using No.2 filter paper and blotting sensor of the Leica GP2 or Vitrobot Mark IV System, respectively. Samples were stored under liquid nitrogen conditions until the time of imaging.

### Cryo-Fluorescence Widefield Microscopy

Fluorescence images of vitrified samples were captured using a THUNDER Imager EM Cryo CLEM widefield microscope (Leica Microsystems), which was equipped with a 50×/0.9 NA dry objective lens, a metal halide light source (EL6000), an air-cooled detector (DFC9000 GT), and a cryo-stage maintained at −190°C. Before observation, grids were placed into AutoGrids™ (Thermo Fisher Scientific). A 2 mm by 2 mm square area was imaged at the grid’s center using the LAS X Navigator software (Leica Microsystems). For each field of view, a symmetrical 20 µm Z-stack with 5 µm intervals was captured centered around the autofocus point to create maps. Subsequently, lamellipodia-forming cells were visually selected for further imaging. A 20 µm Z-stack with 0.75 µm intervals was acquired at these locations. The imaging utilized multiple channels: transmitted light brightfield, reflection, GFP (Ex: BP470/40, Em: 525/50), and Texas Red (Ex: BP560/40, Em: 630/75). Image stitching was executed using the LAS X software. Small Volume Computational Clearing (SVCC) was applied to the acquired image stacks to diminish blurring and enhance weaker or subdued signals. Images and mosaic tiles were exported in TIFF format. Additional image processing, including maximum intensity projection, flipping, cropping, and contrast adjustment, was conducted using ImageJ/Fiji and Adobe Photoshop.

### Cryo-electron microscopy

Cryo-electron tomograms were acquired on a Titan Krios (Thermo Fisher Scientific) operating at an acceleration voltage of 300 kV and equipped with a Cs corrector (CEOS, GmbH), a Volta phase plate and BioQuantam K3 direct detector with energy filter (slit width of 20 eV) (Gatan). The microscope was controlled using the Tomography 5 software (Thermo Fisher Scientific). Initially, an atlas was acquired to identify the square where the target cells were located. Subsequently, the tomography acquisition position was determined based on the correlation with fluorescence images. Tilt series were acquired with a range of up to −70 to 70° in 2° increments. The magnification was ×19,500, resulting in a pixel size of 3.71Å. The electron dose per image was calculated as 1.44 e^-^/Å^2^, with the total dose for series approximately 100 e^-^/Å^2^.

### Image processing

Before 3D reconstruction, poor quality tilt images caused by obstacles such as grid bars blocking the beam at high tilt angles, were removed. The tilt series images were then reconstructed into 3D tomograms by using the IMOD software package^64^. Micrographs were aligned by cross-correlation, followed by alignment through tracking 20 nm gold fiducial beads coated on the grid. These aligned micrographs underwent reconstruction for visual analysis using IMOD SIRT (simultaneous iterative reconstruction technique, number of iterations: 8) at bin2. CTF correction was not performed on these reconstructions.

Segmentation of cellular components was initially performed using AMIRA software (Thermo Fisher Scientific). Before segmentation, the reconstructed images were binned 2 to 3 times (resulting in final bin4 and bin6 from the original images) and processed with a Gaussian filter. The membrane structure was manually segmented, while actin filaments and microtubules were segmented using the XFiber module of Amira^65^. Filament tracing was undertaken after high-contrast structures such as the membrane, grid edge, and outer cellular space were masked. Microtubule tracing utilized the following parameters, as described previously^66^: cylinder length: 600 Å, angular sampling: 5, mask cylinder radius: 140 Å, outer cylinder radius: 125 Å, inner cylinder radius: 75 Å, and missing wedge according to the individual tilt series. For actin filaments, these parameters were used: cylinder length: 500 Å, angular sampling: 6 Å, mask cylinder radius: 45 Å, outer cylinder radius: 35 Å, inner cylinder radius: 0 Å. The Trace Correlation Lines module was configured with these parameter values: minimum seed correlation: 75–125 (tomogram dependent), minimum continuation quality: 60–90 (tomogram dependent), direction coefficient: 0.3, minimum distance: 70 Å, minimum length: 350 Å, search corn minimum step size (%): 10. Segments and point coordinates were extracted into separate Excel sheets from Amira and reformatted to IMOD/Etomo-style files using a MATLAB-script (“amira_reformat_to_coordinates.m” in “computational toolbox for ultrastructural quantitative analysis of filament networks in cryo-ET data”^36^). Tracing errors were manually corrected in 3dmod in IMOD, and the branch points of actin filaments were also determined manually in 3dmod. Quantitative analysis of actin filaments was conducted using the ultrastructural analysis toolbox^36^.

Visualization of the 3D model was performed in 3dmod. Color-coded maps illustrating the angular distribution of actin filaments relative to the leading edge direction were generated using the ultrastructural analysis toolbox. Three-dimensional images of membranes and microtubules images created in 3dmod were meticulously merged with the color maps in Adobe Photoshop.

The distance between the membrane and the tips of actin filaments was measured using ImageJ/Fiji. Graphs were created using Graphpad Prism 10 (GraphPad Software) and MATLAB (MathWorks).

## Data availability

The reconstructed tomograms shown in Figs. 3–5 have been deposited in the EMDB under the following IDs: EMD-61088, EMD-61091, EMD-61092, and EMD-61093. The raw tilt images, prior to 3D reconstruction, as well as the segmentation data has be deposited in the EMPIAR (ID: EMPIAR-12292).

## Acknowledgments

The authors are grateful to Dr. Florian Schur (Institute of Science and Technology (IST), Austria) for helpful discussion about analyzing actin network and 3D printing and Yuriko Sakamaki (the Research Core Center, Institute of Science Tokyo) for technical help. We thank Drs T. Ishii and T. Asano (Institute of Science Tokyo), S. Kikkawa (Kobe University) for the helpful discussion, and S. Nakamura (Institute of Science Tokyo), K. Chin, Y. Sakihama, T. Shimizu (Kobe University), N. Makuta (Osaka University), and M. Nakamura (Mie University) for assistance, and other colleagues in Nakata lab and Nitta lab, the Research Core Center (Institute of Science Tokyo) for usage of the transmission electron microscopy and the carbon coater. This work was partly supported by the Nanotechnology Platform Program and “Advanced Research Infrastructure for Materials and Nanotechnology in Japan (ARIM)” of the Ministry of Education, Culture, Sports, Science and Technology (MEXT, Japan). Proposal Numbers JPMX09A18OS0055, JPMX09A19OS0046, JPMX09A20OS0002, JPMX09A21OS0020, and JPMX12220S0015. This research was also partially supported by the Research Support Project for Life Science and Drug Discovery (Basis for Supporting Innovative Drug Discovery and Life Science (BINDS)) from AMED under Grant Number JP22am121001.

## Author contributions

Conceptualization: H.I., T.I., R.N., T.N.; Methodology: H.I., T.I., A.K., H.T., T.K., K.M., R.T., T.K.; Validation: H.I., T.I., R.N., T.N.; Investigation: H.I., T.I., A.K., S.Y., R.N., T.N.; Resources: H.I., T.I., T.K., H.G., K.M, R.N., T.N.; Data curation: H.I., T.I., K.A., H.T., T.K., M.R., R.N., T.N.; Writing – original draft: H.I.; Writing - review & editing: T.I., S.Y., H.T., R.N., T.N; Visualization: H.I., T.I.; Supervision: R.N., T.N.; Project administration: H.I., T.I., H.G., R.N., T.N.; Funding acquisition: H.I., T.I., S.Y., H.Y., H.T., T.K., H.G., R.N., T.N.

## Abbreviations

Cryo-EM: cryogenic transmission microscopy

Cryo-ET: cryogenic electron tomography

CLEM: correlative light and electron microscopy

ECM: extracellular matrix

EM: electron microscopy

PA-Rac1: photoactivatable Rac1

LM: light microscopy

LLPS: liquid-liquid phase separation

MTs: microtubules

## References

1. 1. Small, J. V., Stradal, T., Vignal, E. & Rottner, K. The lamellipodium: Where motility begins. Trends in Cell Biology vol. 12 Preprint at 10.1016/S0962-8924(01)02237-1 (2002).

2. Innocenti, M. New insights into the formation and the function of lamellipodia and ruffles in mesenchymal cell migration. Cell Adh Migr 12, (2018).

3. Nakata, T. & Hirokawa, N. Cytoskeletal reorganization of human platelets after stimulation revealed by the quick-freeze deep-etch technique. J Cell Biol 105, (1987).

4. Rottner, K., Faix, J., Bogdan, S., Linder, S. & Kerkhoff, E. Actin assembly mechanisms at a glance. J Cell Sci 130, (2017).

5. Svitkina, T. The actin cytoskeleton and actin-based motility. Cold Spring Harb Perspect Biol 10, (2018).

6. Luster, A. D., Alon, R. & von Andrian, U. H. Immune cell migration in inflammation: Present and future therapeutic targets. Nature Immunology vol. 6 Preprint at 10.1038/ni1275 (2005).

7. Charpentier, J. C. & King, P. D. Mechanisms and functions of endocytosis in T cells. Cell Communication and Signaling vol. 19 Preprint at 10.1186/s12964-021-00766-3 (2021).

8. Friedl, P. & Gilmour, D. Collective cell migration in morphogenesis, regeneration and cancer. Nature Reviews Molecular Cell Biology vol. 10 Preprint at 10.1038/nrm2720 (2009).

9. Machesky, L. M. Lamellipodia and filopodia in metastasis and invasion. FEBS Letters vol. 582 Preprint at 10.1016/j.febslet.2008.03.039 (2008).

10. Ridley, A. J., Paterson, H. F., Johnston, C. L., Diekmann, D. & Hall, A. The small GTP-binding protein rac regulates growth factor-induced membrane ruffling. Cell 70, (1992).

11. Nobes, C. D. & Hall, A. Rho, Rac, and Cdc42 GTPases regulate the assembly of multimolecular focal complexes associated with actin stress fibers, lamellipodia, and filopodia. Cell 81, (1995).

12. Miki, H., Suetsugu, S. & Takenawa, T. WAVE, a novel WASP-family protein involved in actin reorganization induced by Rac. EMBO Journal 17, (1998).

13. Machesky, L. M. et al. Scar, a WASp-related protein, activates nucleation of actin filaments by the Arp2/3 complex. Proc Natl Acad Sci U S A 96, (1999).

14. Mullins, R. D., Heuser, J. A. & Pollard, T. D. The interaction of Arp2/3 complex with actin: Nucleation, high affinity pointed end capping, and formation of branching networks of filaments. Proc Natl Acad Sci U S A 95, (1998).

15. Mattila, P. K. & Lappalainen, P. Filopodia: Molecular architecture and cellular functions. Nature Reviews Molecular Cell Biology vol. 9 Preprint at 10.1038/nrm2406 (2008).

16. Watanabe, N. & Mitchison, T. J. Single-molecule speckle analysis of actin filament turnover in lamellipodia. Science (1979) 295, (2002).

17. Yamashiro, S. et al. New single-molecule speckle microscopy reveals modification of the retrograde actin flow by focal adhesions at nanometer scales. Mol Biol Cell 25, (2014).

18. Mehidi, A. et al. Forces generated by lamellipodial actin filament elongation regulate the WAVE complex during cell migration. Nat Cell Biol 23, (2021).

19. Rimoli, C. V., Valades-Cruz, C. A., Curcio, V., Mavrakis, M. & Brasselet, S. 4polar-STORM polarized super-resolution imaging of actin filament organization in cells. Nat Commun 13, (2022).

20. Wu, C. et al. Arp2/3 is critical for lamellipodia and response to extracellular matrix cues but is dispensable for chemotaxis. Cell 148, (2012).

21. Ponti, A. et al. Periodic patterns of actin turnover in lamellipodia and lamellae of migrating epithelial cells analyzed by quantitative fluorescent speckle microscopy. Biophys J 89, (2005).

22. Koestler, S. A., Auinger, S., Vinzenz, M., Rottner, K. & Small, J. V. Differentially oriented populations of actin filaments generated in lamellipodia collaborate in pushing and pausing at the cell front. Nat Cell Biol 10, (2008).

23. Svitkina, T. M. & Borisy, G. G. Arp2/3 complex and actin depolymerizing factor/cofilin in dendritic organization and treadmilling of actin filament array in lamellipodia. Journal of Cell Biology 145, (1999).

24. Svitkina, T. M., Verkhovsky, A. B., McQuade, K. M. & Borisy, G. G. Analysis of the actin-myosin II system in fish epidermal keratocytes: Mechanism of cell body translocation. Journal of Cell Biology 139, (1997).

25. Schneider, J. & Jasnin, M. Capturing actin assemblies in cells using in situ cryo-electron tomography. Eur J Cell Biol 101, (2022).

26. Young, L. N. & Villa, E. Bringing Structure to Cell Biology with Cryo-Electron Tomography. Annual Review of Biophysics vol. 52 Preprint at 10.1146/annurev-biophys-111622-091327 (2023).

27. Fäßler, F., Javoor, M. G. & Schur, F. K. M. Deciphering the molecular mechanisms of actin cytoskeleton regulation in cell migration using cryo-EM. Biochemical Society Transactions vol. 51 Preprint at 10.1042/BST20220221 (2023).

28. Vinzenz, M. et al. Actin branching in the initiation and maintenance of lamellipodia. J Cell Sci 125, (2012).

29. Mueller, J. et al. Load Adaptation of Lamellipodial Actin Networks. Cell 171, (2017).

30. Fäßler, F., Dimchev, G., Hodirnau, V. V., Wan, W. & Schur, F. K. M. Cryo-electron tomography structure of Arp2/3 complex in cells reveals new insights into the branch junction. Nat Commun 11, (2020).

31. 31. Chung, W. L., et al. A network of mixed actin polarity in the leading edge of spreading cells. Commun Biol 5, (2022).

32. Wu, Y. I. et al. A genetically encoded photoactivatable Rac controls the motility of living cells. Nature 461, 104–108 (2009).

33. Riedl, J. et al. Lifeact: A versatile marker to visualize F-actin. Nat Methods 5, (2008).

34. Nedozralova, H. et al. In situ cryo-electron tomography reveals local cellular machineries for axon branch development. Journal of Cell Biology 221, (2022).

35. Aramaki, S., Mayanagi, K., Jin, M., Aoyama, K. & Yasunaga, T. Filopodia formation by crosslinking of F-actin with fascin in two different binding manners. Cytoskeleton 73, (2016).

36. Dimchev, G., Amiri, B., Fäßler, F., Falcke, M. & Schur, F. K. Computational toolbox for ultrastructural quantitative analysis of filament networks in cryo-ET data. J Struct Biol 213, (2021).

37. Jansen, S. et al. Mechanism of actin filament bundling by fascin. Journal of Biological Chemistry 286, (2011).

38. Yang, S. et al. Molecular mechanism of fascin function in filopodial formation. Journal of Biological Chemistry 288, (2013).

39. Langanger, G. et al. Ultrastructural localization of α-actinin and filamin in cultured cells with the immunogold staining (IGS) method. Journal of Cell Biology 99, (1984).

40. Salmon, W. C., Adams, M. C. & Waterman-Storer, C. M. Dual-wavelength fluorescent speckle microscopy reveals coupling of microtubule and actin movements in migrating cells. Journal of Cell Biology 158, (2002).

41. Gupton, S. L., Salmon, W. C. & Waterman-Storer, C. M. Converging populations of f-actin promote breakage of associated microtubules to spatially regulate microtubule turnover in migrating cells. Current Biology 12, (2002).

42. Küppers, M., Albrecht, D., Kashkanova, A. D., Lühr, J. & Sandoghdar, V. Confocal interferometric scattering microscopy reveals 3D nanoscopic structure and dynamics in live cells. Nat Commun 14, (2023).

43. Urban, E., Jacob, S., Nemethova, M., Resch, G. P. & Small, J. V. Electron tomography reveals unbranched networks of actin filaments in lamellipodia. Nat Cell Biol 12, (2010).

44. Boujemaa-Paterski, R. et al. Network heterogeneity regulates steering in actin-based motility. Nat Commun 8, (2017).

45. Suraneni, P. et al. The Arp2/3 complex is required for lamellipodia extension and directional fibroblast cell migration. Journal of Cell Biology 197, (2012).

46. Case, L. B., Zhang, X., Ditlev, J. A. & Rosen, M. K. Stoichiometry controls activity of phase-separated clusters of actin signaling proteins. Science (1979) 363, (2019).

47. Yang, S. et al. Self-construction of actin networks through phase separation-induced abLIM1 condensates. Proc Natl Acad Sci U S A 119, (2022).

48. Graham, K. et al. Liquid-like VASP condensates drive actin polymerization and dynamic bundling. Nat Phys 19, (2023).

49. 49. Faix, J. & Rottner, K. Ena/VASP proteins in cell edge protrusion, migration and adhesion. Journal of Cell Science vol. 135 Preprint at 10.1242/jcs.259226 (2022).

50. Harker, A. J. et al. Ena/VASP processive elongation is modulated by avidity on actin filaments bundled by the filopodia cross-linker fascin. Mol Biol Cell 30, (2019).

51. Yamashiro, S. & Watanabe, N. A new link between the retrograde actin flow and focal adhesions. Journal of Biochemistry vol. 156 Preprint at 10.1093/jb/mvu053 (2014).

52. Peskin, C. S., Odell, G. M. & Oster, G. F. Cellular motions and thermal fluctuations: the Brownian ratchet. Biophys J 65, (1993).

53. Vallotton, P., Gupton, S. L., Waterman-Storert, C. M. & Danuser, G. Simultaneous mapping of filamentous actin flow and turnover in migrating cells by quantitative fluorescent speckle microscopy. Proc Natl Acad Sci U S A 101, (2004).

54. Yamashiro, S. et al. Convection-Induced Biased Distribution of Actin Probes in Live Cells. Biophys J 116, (2019).

55. Shaaya, M., Fauser, J. & Karginov, A. V. Optogenetics: The art of illuminating complex signaling pathways. Physiology 36, (2021).

56. Tischer, D. & Weiner, O. D. Illuminating cell signalling with optogenetic tools. Nat Rev Mol Cell Biol 15, 551–558 (2014).

57. Chakrabarti, R. et al. Optogenetics and electron tomography for structure-function analysis of cochlear ribbon synapses. Elife 11, (2022).

58. Watanabe, S. Flash-and-freeze: Coordinating optogenetic stimulation with rapid freezing to visualize membrane dynamics at synapses with millisecond resolution. Front Synaptic Neurosci 8, (2016).

59. Romero, A. M., Bonin, C. & Twomey, E. C. Time-Resolved Cryo-EM Specimen Preparation with Single Millisecond Precision. bioRxiv 2023.08.24.554704 (2023) doi:10.1101/2023.08.24.554704.

60. Fäßler, F., Zens, B., Hauschild, R. & Schur, F. K. M. 3D printed cell culture grid holders for improved cellular specimen preparation in cryo-electron microscopy. J Struct Biol 212, (2020).

61. Kakumoto, T. & Nakata, T. Optogenetic Control of PIP3: PIP3 Is Sufficient to Induce the Actin-Based Active Part of Growth Cones and Is Regulated via Endocytosis. PLoS One 8, 1–17 (2013).

62. Inaba, H., Miao, Q. & Nakata, T. Optogenetic control of small GTPases reveals RhoA mediates intracellular calcium signaling. Journal of Biological Chemistry 100290 (2021) doi:10.1016/j.jbc.2021.100290.

63. 63. Schindelin, J., et al. Fiji: An open-source platform for biological-image analysis. Nature Methods Preprint at 10.1038/nmeth.2019 (2012).

64. Kremer, J. R., Mastronarde, D. N. & McIntosh, J. R. Computer visualization of three-dimensional image data using IMOD. J Struct Biol 116, (1996).

65. Rigort, A. et al. Automated segmentation of electron tomograms for a quantitative description of actin filament networks. J Struct Biol 177, (2012).

66. Watanabe, R. et al. The In Situ Structure of Parkinson’s Disease-Linked LRRK2. Cell 182, (2020).

