## Supplementary Information for "Cryo-ET of actin cytoskeleton and membrane structure in lamellipodia formation using optogenetics"

**Inaba et al., Fig. S1**

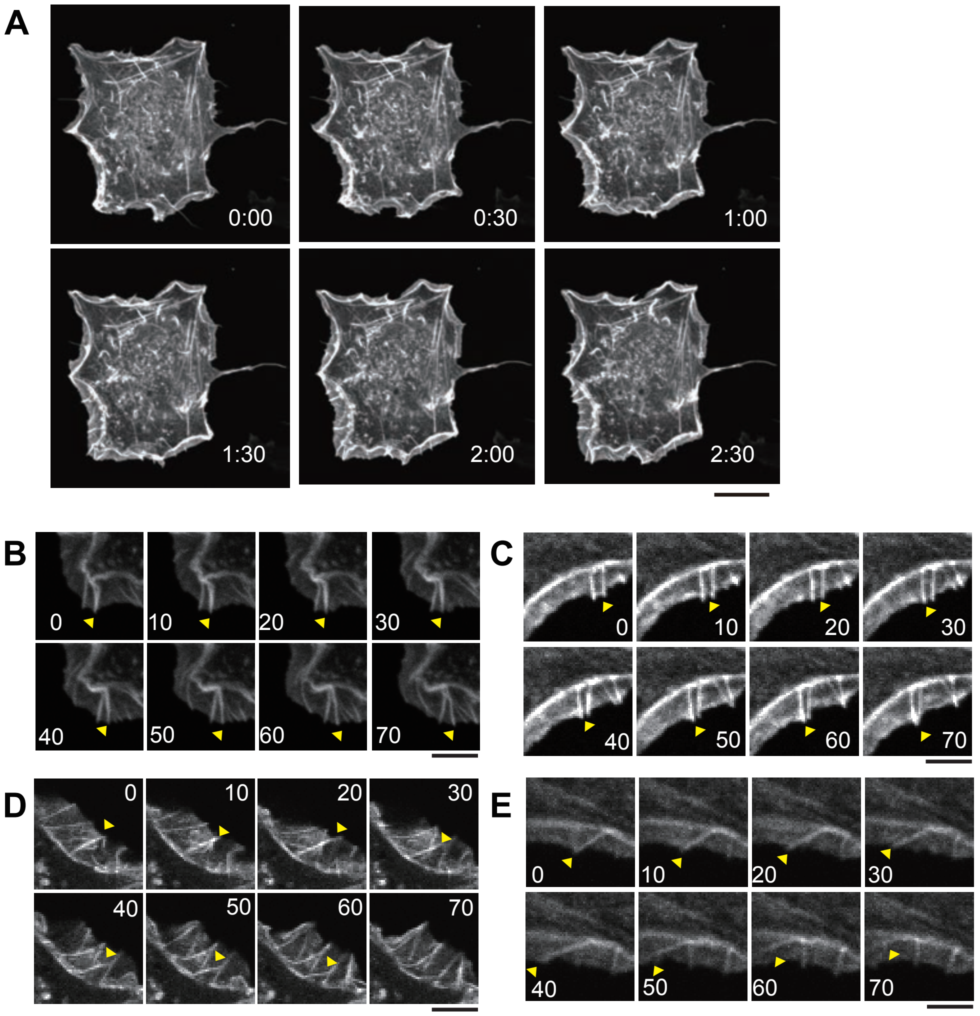


**Fig. S1. PA-Rac1-induced lamellipodia formation and its dynamics.** COS-7 cells expressing PA-Rac1 and Lifeact-mCherry on glass bottom dishes **(A, B, D)** or EM grids **(C, E)**. Images were captured every 10 sec, with PA-Rac1 activated by a 458 nm laser from time 0 at 10-sec intervals. The fluorescence images of Lifeact-mCherry are shown. **(A)** Representative time-lapse images on glass-bottom dishes. **(B, C)** Detail the bilateral movement of microspikes and filopodia within lamellipodia on glass-bottom dishes **(B)** and EM grids **(C)**. **(D, E)** Collapse of microspikes or filopodia in cells on glass-bottom **(D)** and EM grids **(E)**. The numbers in each panel indicate time in seconds, and yellow arrowheads indicate the microspikes and filopodia in **B–D**. Scale bar = 20 μm in **A** and 5 μm in **B–E**, respectively.

Inaba et al., Fig. S2

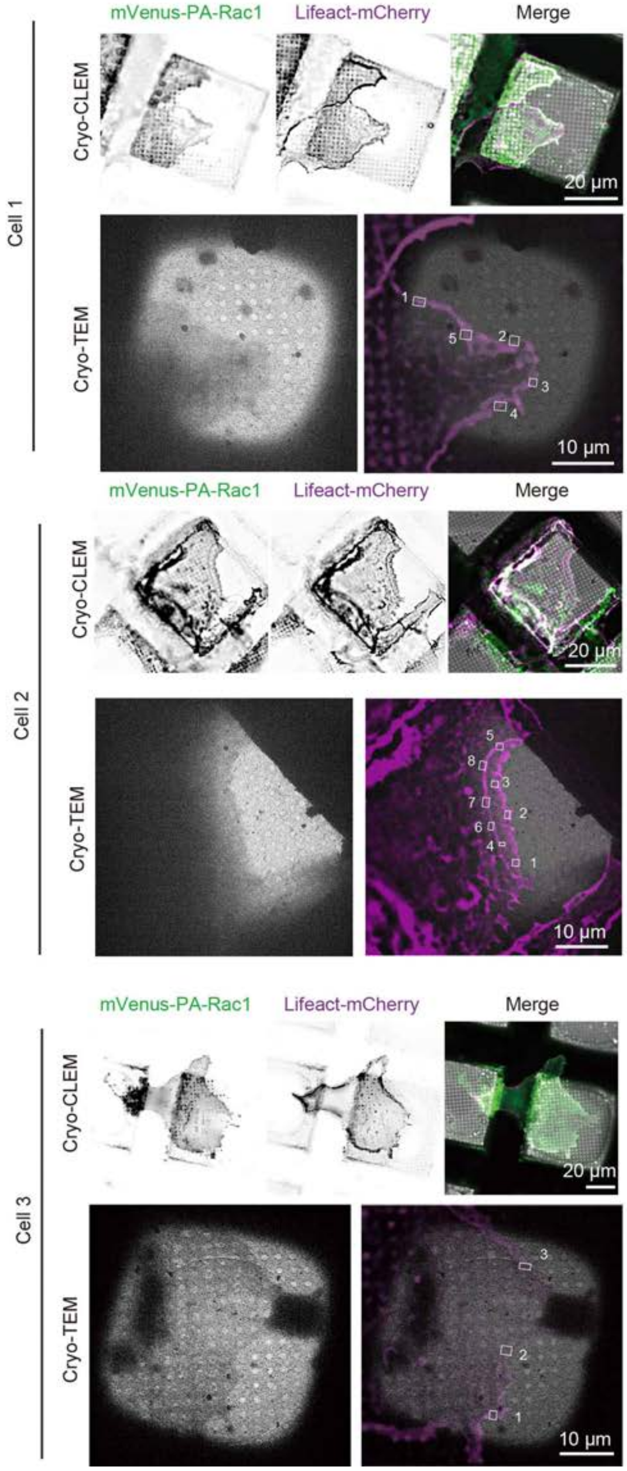


**Fig. S2. Cryo-CLEM images.** Cryo-CLEM images of three cells which tomograms were captured in this study. Green: mVenus-PA-Rac1, Purple: Lifeact-mCherry.

**Inaba et al., Fig. S3**

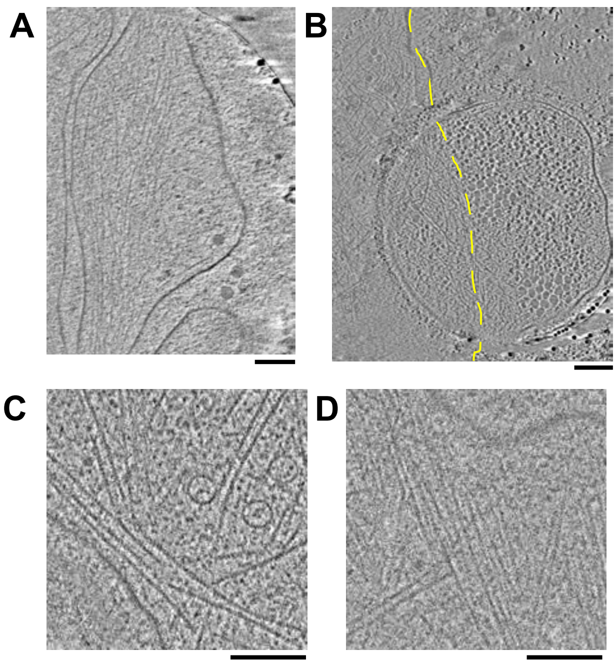


**Fig. S3. Deformation of the plasma membrane at the hole in Quantifoil grids. (A, B)** Deformation of the plasma membrane at filopodia **(A)** and lamellipodia **(B)** at Quantifoil's hole. In **A**, actin bundles extend through the center, while the plasma membrane exhibits a bulge to the right, containing few filaments within. In **B**, the yellow dashed line indicates the presumed leading edge from the cell membrane outside the hole. However, inside the hole, the plasma membrane bulges towards the right, containing few filaments. **(C, D)** Contrast differences in images from the same tomography captured inside **(C)** and outside **(D)** the hole. Scale bar = 200 nm.

Inaba et al., Fig. S4

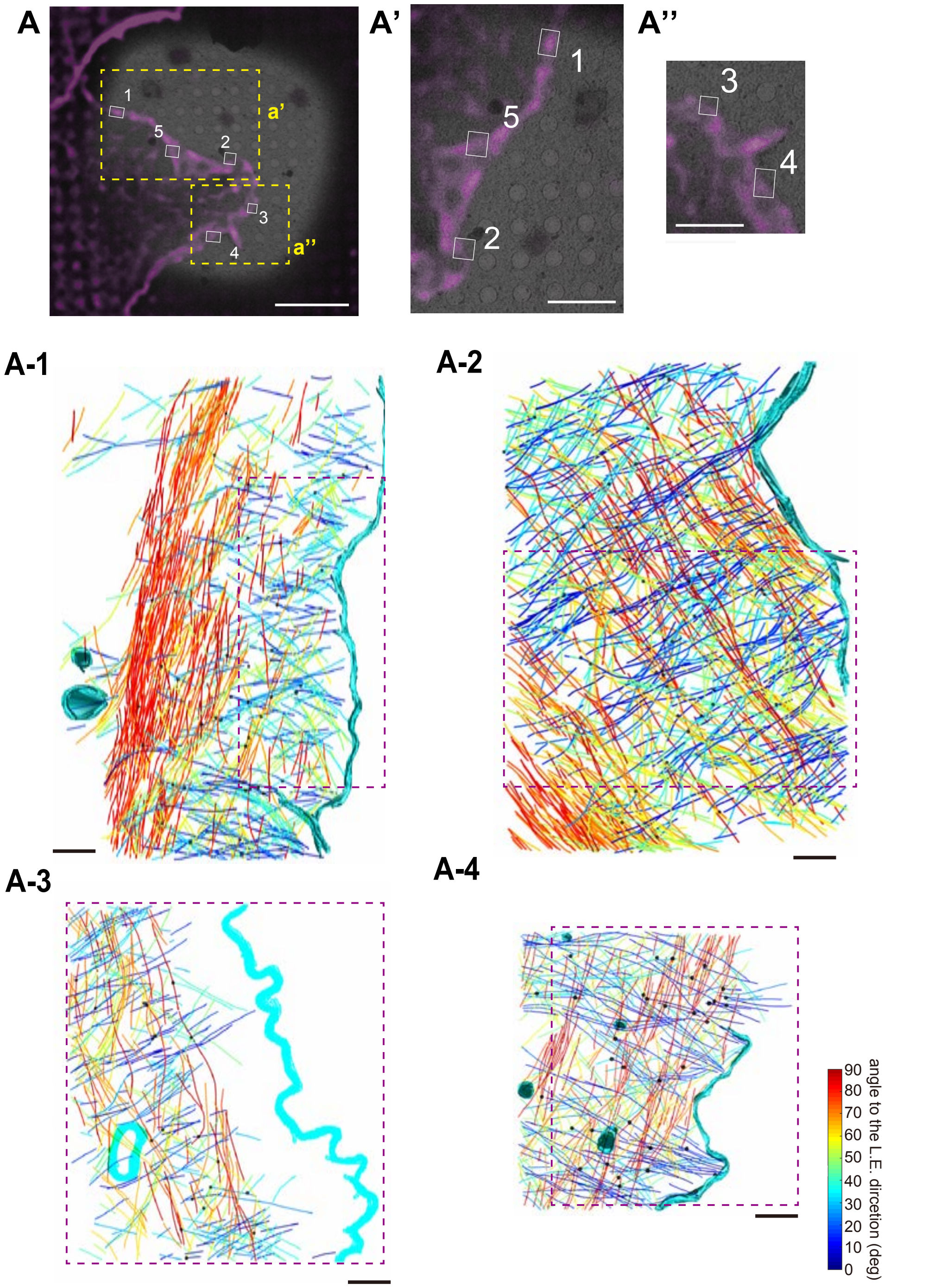


**Fig. S4. Gallery of Cryo-CLEM and segmented images from cryo-ET.** White boxes in the CLEM images indicate the areas where cryo-ET was acquired. In the segmented images, the plasma membrane is represented in cyan microtubules in green, and ER in blue. Actin filaments are color-coded to illustrate their orientation relative to the leading edge (L.E.) of the cells. The yellow dotted box in **A**, **B**, and **C** are magnified in **A'** and **A"**, **B'**, and **C'** and **C"**, respectively. The purple dotted box highlights the lamellipodia region. Scale bars: 10 µm in cryo-CLEM images and 200 nm in segmented images.

Inaba et al., Fig. S4

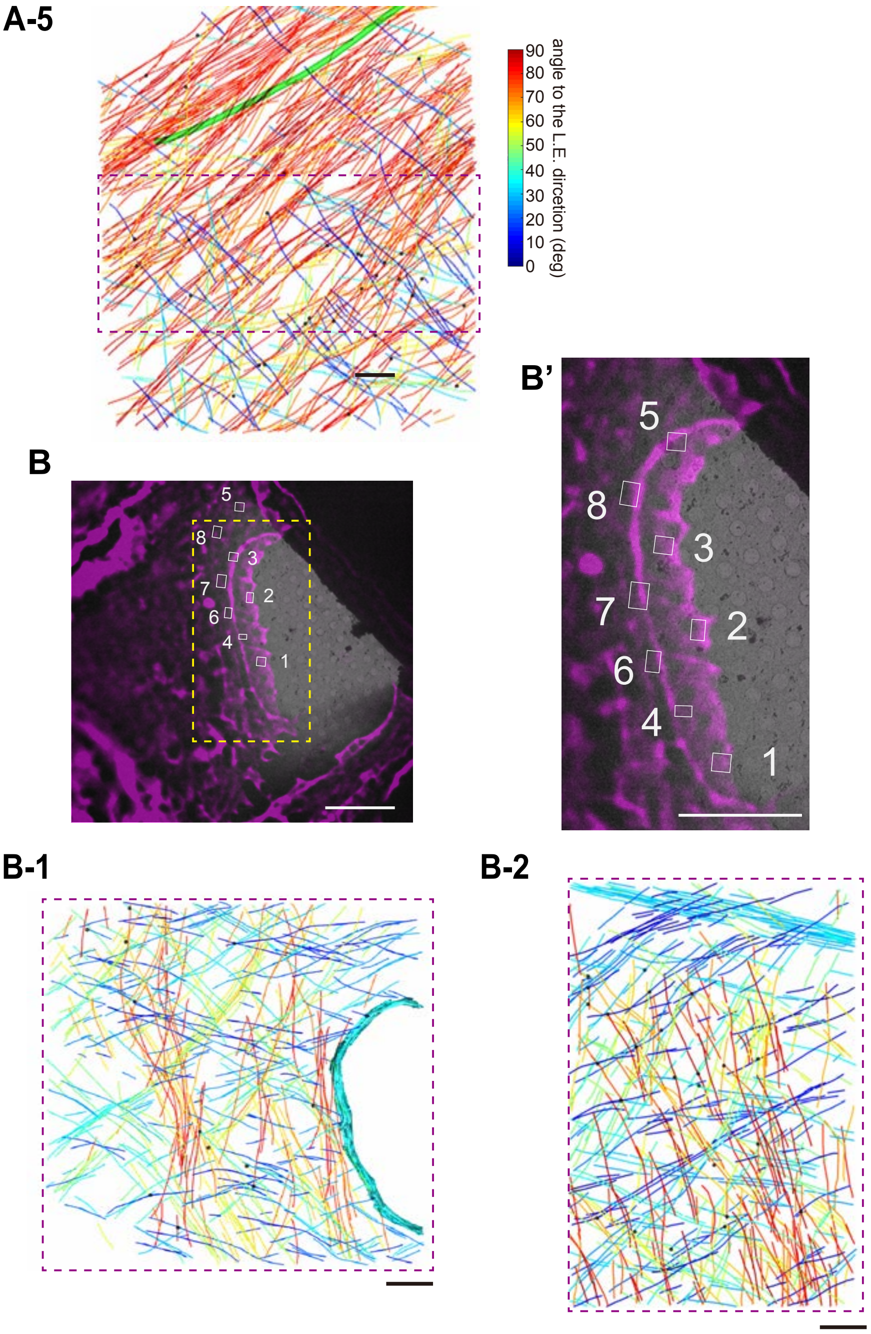


**Fig. S4 Continuation**

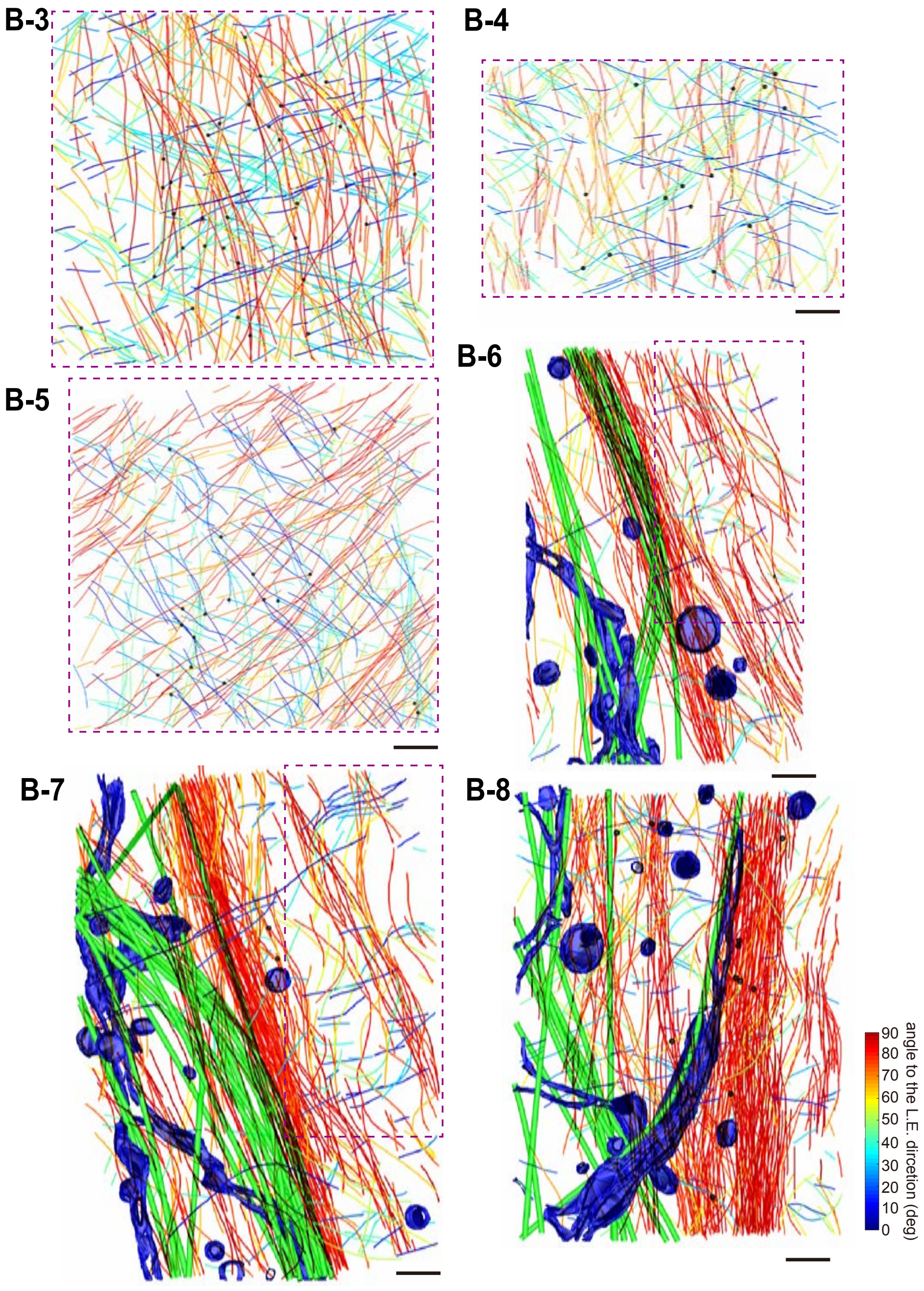


**Fig. S4 Continuation**

Inaba et al., Fig. S4

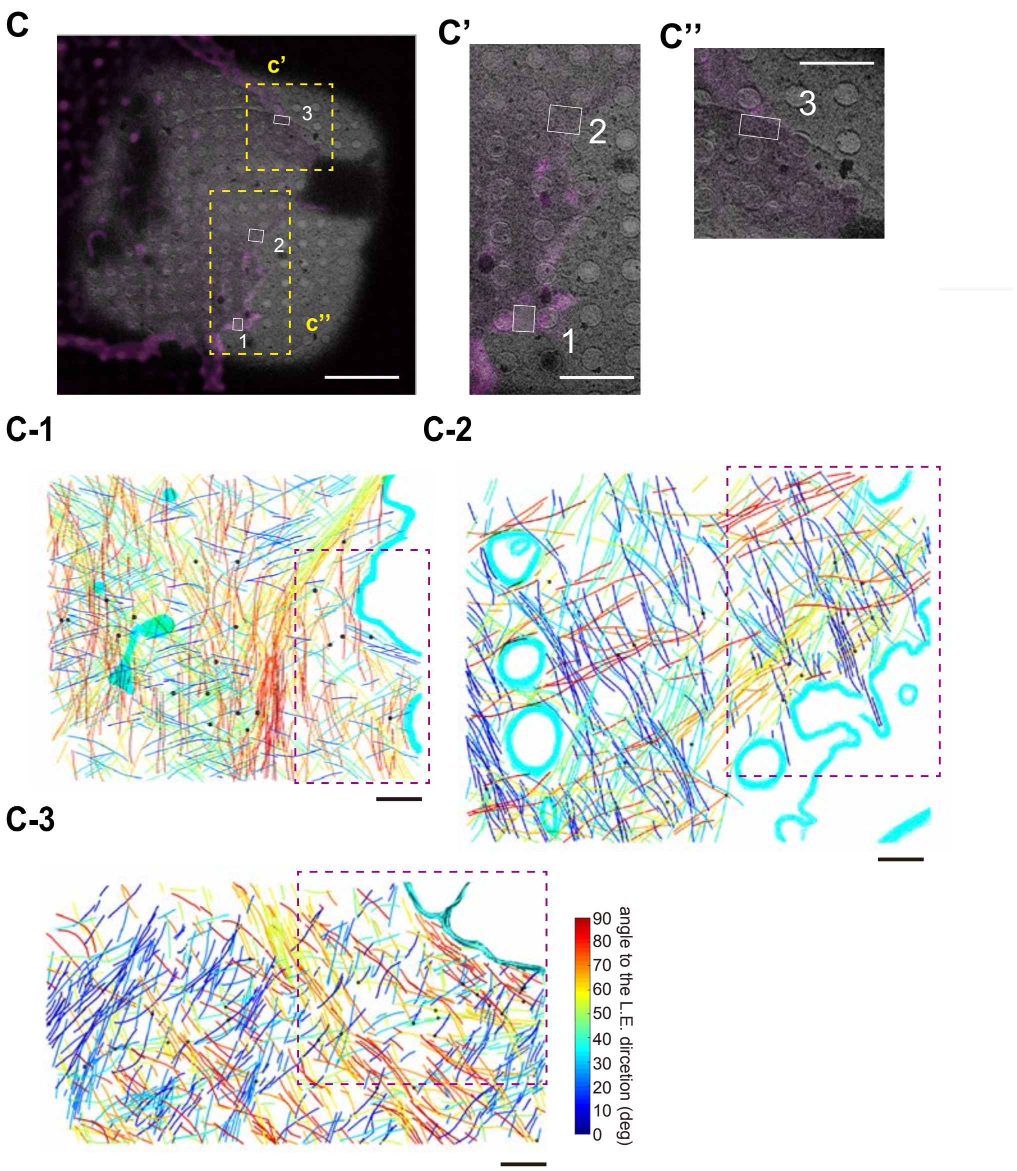


**Fig. S4 Continuation**

**Inaba et al., Fig. S5**

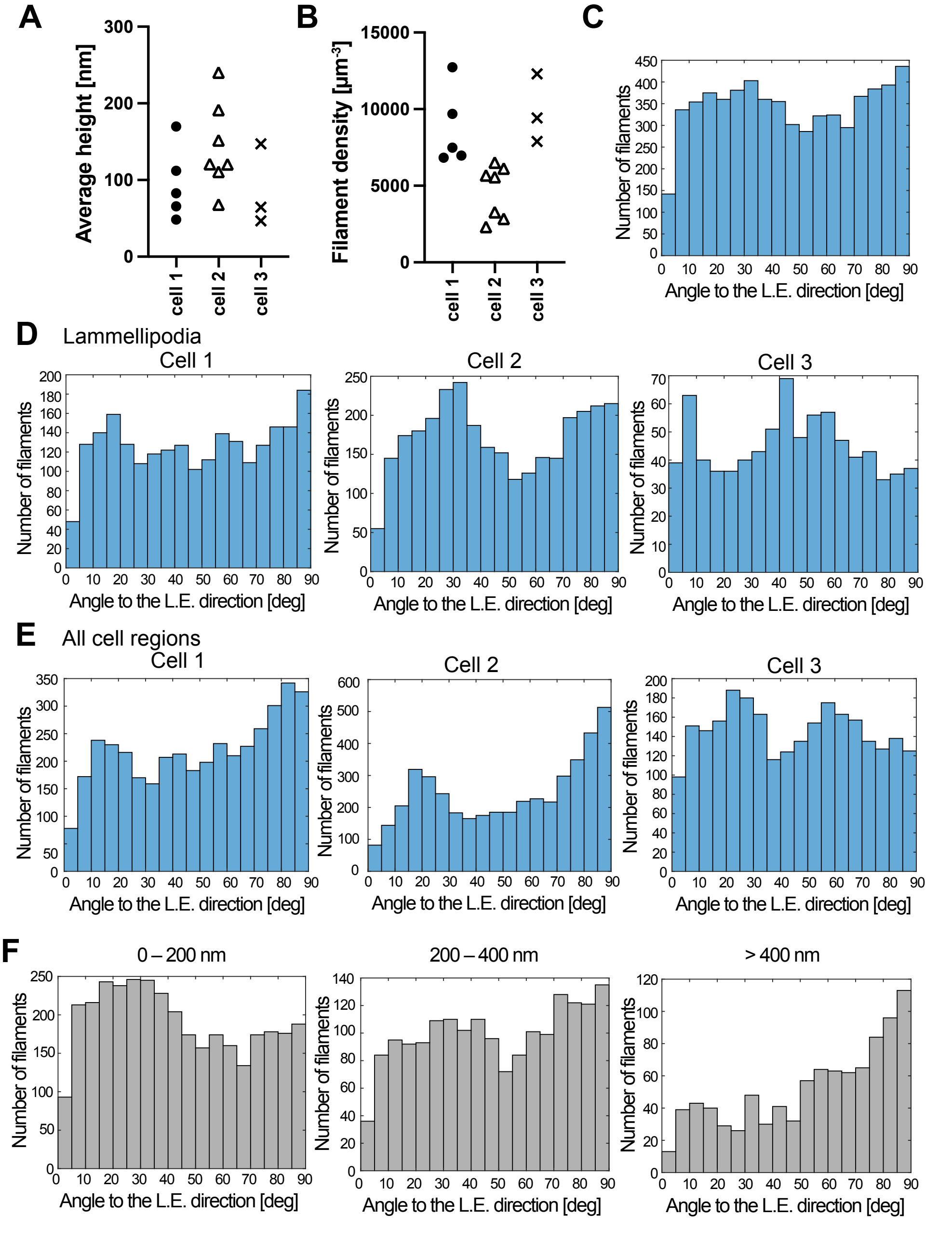


**Fig. S5. The organization of the actin cytoskeleton in PA-Rac1 induced lamellipodia (A, B)** The averaged height **(A)** and the filament density **(B)** in lamellipodia across individual cryo-ET. **(C–E)** The orientation of the actin filaments relative to the leading edge (L.E.) in lamellipodia (**C, D**) and across all cellular regions (**E**) from three cells (**C**) and individual cells (**D, E**). **(F)** Orientation distributions of actin filaments by length: 0–200 nm (left), 200–400 nm (center), >400 nm (right).

Inaba et al., Fig. S6

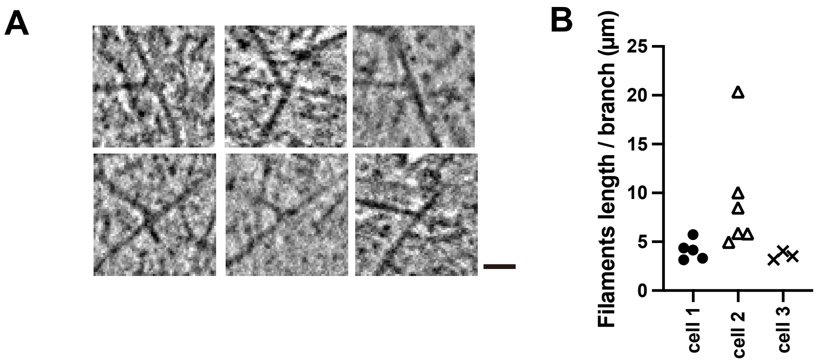


**Fig. S6. Actin branches within lamellipodia.** (**A**) Representative images of branched actin filaments. Scale bar = 25 nm. (**B**) The frequency of actin branches in lamellipodia across individual cryo-ET.

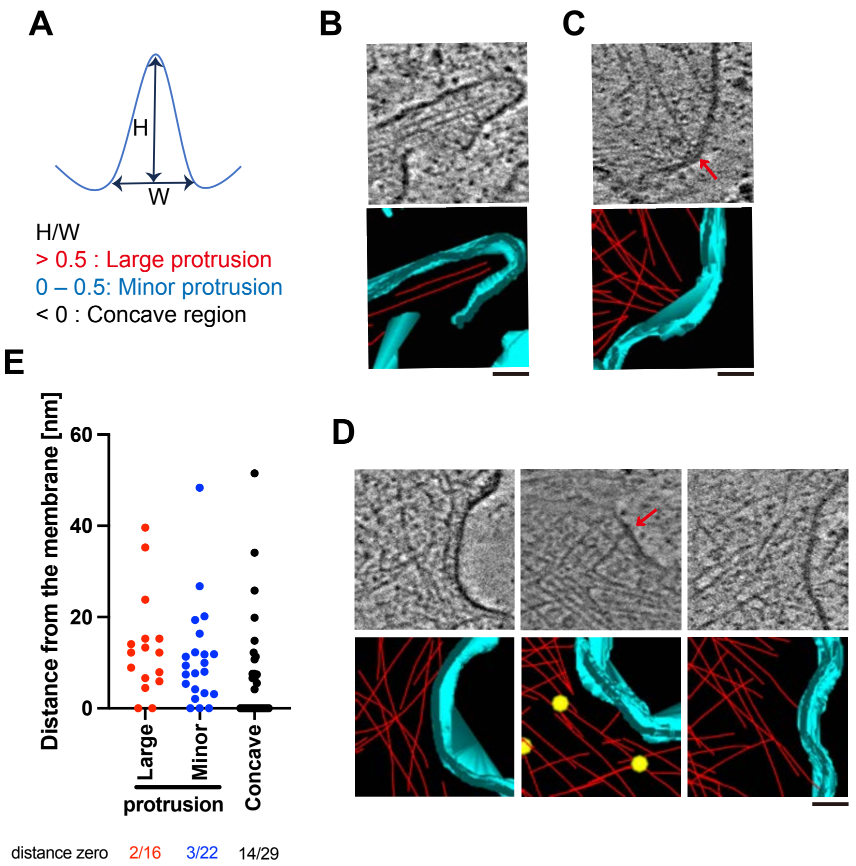


**Figure S7**. **The architecture of the plasma membrane at the leading edge and the associated actin network. (A)** A schematic representation of various membrane architectures. Membrane protrusions are measured by their height ("H") and base width ("W"). They are categorized by the W/H ratio: "Large protrusion" for W/H over 0.5, "Minor protrusion" for W/H between 0 and 0.5 and "Concave regions" for W/H less than 0. **(B–D)** Gallery views of each protrusion type: "Large protrusion" **(B)**, "Minor protrusion" **(C)**, and "Concave regions" **(D)**. Yellow dots mark branched points of actin filaments; red arrows point to locations where actin filaments attach to the plasma membrane; an orange asterisk denotes regions where the plasma membrane shows high intensity. Scale bar = 50 nm. **(E)** Analysis of the distance between the membrane and the ends of actin filaments in the three types of membrane architectures. Only actin filaments oriented towards the leading edge were analyzed. The number of direct interaction points, relative to the total analyzed actin filaments, is noted below the graph.

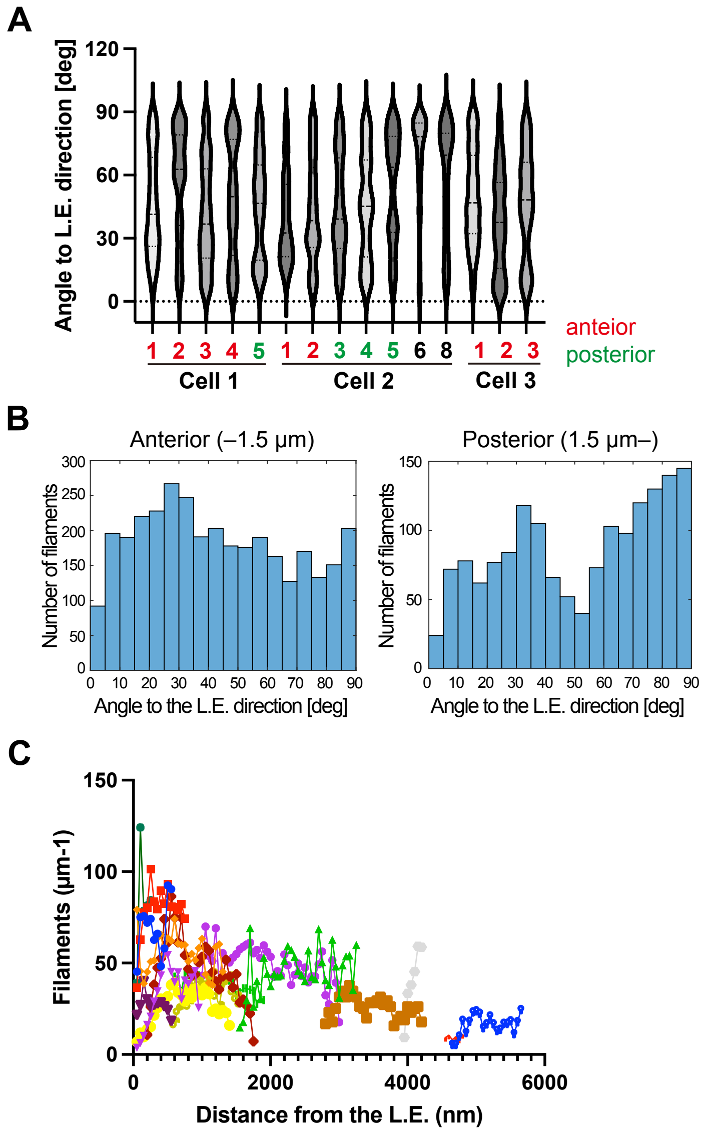


**Fig. S8. The orientation and the density of actin filaments relative to the leading edge (L.E.)** (**A**) The orientation of actin filaments relative to the L.E. Tomography numbers designated as "anterior region" are marked in red, while those identified as "posterior region" are noted in green. **(B)** The orientation distributions of actin filaments in anterior regions (left: approximately 1.5 µm from the L.E.) and posterior region (right: more than 1.5 µm from L.E.). **(C)** The density of actin filaments is relative to the L.E., where straight lines were drawn at 50 nm intervals from the L.E., and the number of intersecting actin filaments was counted. The data obtained from each tomography were plotted on a graph.

**Video 1.** PA-Rac1-induced lamellipodia formation on the glass-bottom dishes. Supplementary movie for Fig. S1.

**Video 2.** PA-Rac1-induced lamellipodia formation on the EM grids. Supplementary movie for Fig. 1D.

**Video 3.** Cryo-ET of PA-Rac1-induced lamellipodia in the early stage of formation. The tomogram corresponds to Fig. 3D. Red: actin, Cyan: membrane, Yellow: actin branches. Scale bar = 100 nm.

**Video 4.** Cryo-ET of PA-Rac1-induced lamellipodia in the middle stage of formation. The tomogram corresponds to Fig. 4D. Scale bar = 200 nm.

**Video 5.** Cryo-ET of PA-Rac1-induced lamellipodia in the late stage of formation. The tomogram corresponds to Fig. 5D. Scale bar = 200 nm.

**Video 6.** Cryo-ET of PA-Rac1-induced lamellipodia in the late stage of formation. The tomogram corresponds to Fig. 5F. Scale bar = 200 nm.
